# Minor isozymes tailor yeast metabolism to carbon availability

**DOI:** 10.1101/394056

**Authors:** Patrick H. Bradley, Patrick A. Gibney, David Botstein, Olga G. Troyanskaya, Joshua D. Rabinowitz

## Abstract

Isozymes are enzymes that differ in sequence but catalyze the same chemical reactions. Despite their apparent redundancy, isozymes are often retained over evolutionary time for reasons that can be unclear. We find that, in yeast, isozymes are strongly enriched in central carbon metabolism. Using a gene expression compendium, we find that many isozyme pairs show anticorrelated expression during the respirofermentative shift, suggesting roles in adapting to changing carbon availability. Building on this observation, we assign function to two minor central carbon isozymes, aconitase 2 (*ACO2*) and pyruvate kinase 2 (*PYK2*). *ACO2* is expressed during fermentation and proves advantageous when glucose is limiting. *PYK2* is expressed during respiration and proves advantageous for growth on three-carbon substrates. *PYK2*’s deletion is rescued by expressing the major pyruvate kinase, but only if that enzyme carries mutations mirroring *PYK2*’s allosteric regulation. Thus, central carbon isozymes enable more precise tailoring of metabolism to changing nutrient availability.

**Importance:** Gene duplication is one of the main evolutionary drivers of new protein function. However, some gene duplicates have nevertheless persisted long-term without apparent divergence in biochemical function. Further, under standard lab conditions, many isozymes have subtle or no knockout phenotypes. These factors make it hard to assess the unique contributions of individual isozymes to fitness. We therefore developed a method to identify experimental perturbations that could expose such contributions, and applied it to yeast gene expression data, revealing a potential role for a set of yeast isozymes in adapting to changing carbon sources. Our experimental confirmation of distinct roles for two “minor” yeast isozymes, including one with no previously described knockout phenotype, highlight that even apparently redundant paralogs can have important and unique functions, with implications for genome-scale metabolic modeling and systems-level studies of quantitative genetics.

## Introduction

Isozymes are distinct proteins within a single organism that can catalyze the same biochemical reactions. Although some isozymes differ in localization, substrate specificity, or cofactor preference, there are also many isozymes that are not differentiated by these criteria. The genome of budding yeast (*Saccharomyces cerevisiae*) contains many duplicate genes encoding isozymes that have persisted since the ancient duplication of the whole genome that led to the evolution of the modern *Saccharomyces* (1). Only a small fraction of these yeast gene duplications remain, strongly suggesting that the remaining ones, including those that encode isozymes, must somehow have contributed to evolutionary fitness.

Several explanations, both complementary and also at times conflicting, have been advanced for the retention of such isozymes (and gene duplicates more generally). One is gene dosage, in which multiple gene copies contribute to maintaining adequate total enzyme levels. Papp et al. have argued that many isozyme pairs can be explained by gene dosage, since in a flux-balance model reactions catalyzed by isozymes tended to carry higher flux (2). However, a subsequent study using experimentally determined fluxes estimated that less than 20% of isozyme pairs catalyzed high flux reactions (3). Additionally, in some high-flux reactions, such as aconitase and pyruvate kinase, one “major” isozyme but not the other “minor” isozyme has been found to be essential under laboratory conditions.

Another potential explanation involves genetic backup, i.e., the ability of isozymes to compensate for the deletion of their partners. However, since genetic backup cannot be directly selected, it is generally agreed that this is more likely to be a side effect of isozyme retention than the cause (4). Kafri et al. demonstrated that some isozymes change in expression after deletion of their partners (“transcriptional reprogramming”), and argued that selection for robustness against nongenetic noise could give rise to both transcriptional reprogramming and genetic backup (5, 6); however, a follow-up study reported that transcriptional reprogramming was only confirmed in ∼11% of tested isozyme pairs (7).

Isozymes are often differentially regulated, suggesting a role in fine-tuning metabolic capabilities (8). A well-understood example of such fine-tuning involves the seven hexose transporters of *S. cerevisiae* (*HXT1-7*), some of which are high-affinity/low-flux and others are low-affinity/high-flux. Collectively, these transporters allow yeast to import hexose optimally across a wide variety of environmental conditions (9). Another form of fine-tuning involves optimization for growth under specific (and less commonly-studied) environmental conditions, and it has been argued that isozymes contribute to such optimization (2, 10, 11). However, so far, existing computational and experimental tools have not proven well-suited to finding the most relevant environmental conditions for explaining the existence of isozymes. For example, flux balance analysis (FBA) models metabolism at the level of reactions, not genes, and is therefore intrinsically unable to differentiate between isozymes (2, 12, 13).

High throughput experimental methods are, in principle, well suited to identifying the function of isozymes. Most isozymes have been knocked out in *Saccharomyces cerevisiae*, and the growth rate and competitive fitness of the resulting strains measured (3, 11, 14–16). A large number of isozyme deletions, however, have failed to show substantial fitness defects under laboratory growth conditions. For example, in a recent study that measured competitive fitness to high precision, 65% of the assayed isozyme deletions had relative fitnesses ≥ 0.99 and some “minor” isoforms, such as the pyruvate kinase isozyme *PYK2*, even showed a slight fitness advantage (15). A limitation of these studies is that they have been conducted in only a few environmental conditions, mainly growth on rich media or defined media with amino acids, with glucose (and sometimes ethanol or glycerol) as the carbon source. The genetic tools that enable these massively parallel assays also tend to use amino acid auxotrophies as selectable markers; growth of these knockout strains therefore requires nutritional supplements that can themselves contribute to growth – clearly less than ideal for studying the function of genes in central metabolism, as has indeed been recently demonstrated (17).

In contrast, transcriptional profiling has been conducted in a much wider array of experimental conditions. Thus, an alternative approach to identifying the function of isozymes is to mine compendia of gene expression data, with the aim of identifying conditions under which isozymes may contribute to fitness. Indeed, previous studies have noted that the differential expression of isozymes is a feature of many microarray experiments (18, 19). However, existing expression analyses (5, 19) have tended to focus on identifying transcriptional co-regulation of isozymes with other enzymes or processes. They have not focused on generating hypotheses about which environments are specifically associated with isozyme function.

Here we develop methods for systematically associating isozymes with specific environmental perturbations, and use these methods to identify an important role for isozymes in adapting to changing carbon source availability. This observation is intriguing given that many (though not all) metabolic isozymes date from the events that may have led *Saccharomyces* to adopt a bifurcated lifestyle, primarily fermenting when glucose is present and respiring otherwise (in what is called the “Crabtree effect”) (20–22). It suggests a rationale for the retention of isozymes over evolutionary time, providing flexibility to central metabolism and in particular, central carbon metabolism. We tested for such flexibility experimentally, by growing cells lacking specific isozymes in alternative carbon sources. In two cases we found growth defects for isozyme deletions on non-standard carbon sources, associating for the first time a specific functional role to the genes that encode them. These experimental results support for the idea that central carbon metabolic isozymes have been retained over evolutionary time to optimize the metabolism of diverse carbon sources.

## Results

### Differentially expressed isozymes are prevalent in central carbon metabolism

We began by assembling a list of co-localized metabolic isozymes (see Methods). We found that, like duplicated yeast genes in general (22), these isozymes concentrate in central carbon metabolism (Supplemental Figure S1, Fisher’s test *p*=1.04×10^−9^). Nearly every step in glycolysis and gluconeogenesis can be catalyzed by more than one enzyme, and storage carbohydrate metabolism and the pentose phosphate pathway also contain many isozyme pairs. In contrast, while many metabolic enzymes are involved in amino acid *de novo* biosynthesis, these pathways contain comparatively few isozymes. Indeed, in the Yeast Pathway database (23), 8% (39/485) of reactions overall are catalyzed by isozymes; however, in the pathways of glycolysis, gluconeogenesis, and fermentation, this number rises to 75% (9/12, Bonferroni-Holm-corrected Fisher’s test p=4.7×10^−8^). The pentose phosphate pathway and TCA/glyoxylate cycle are also significantly enriched for reactions catalyzed by isozymes (Bonferroni-Holm corrected Fisher’s test p=0.03 and p=0.01, respectively; see Table S1).

We also note that metabolic isozymes were strongly enriched for genes dating from the whole-genome duplication (WGD) of yeast (63% of isozymes date from the WGD, compared with 19% of the genome; Fisher’s test p < 10^−22^). Compared to non-WGD yeasts, post-WGD yeasts such as *Saccharomyces cerevisiae* are more likely to exhibit the Crabtree effect, i.e., to ferment glucose to ethanol even in the presence of oxygen. In addition, post-WGD yeast are more likely to be able to survive without the mitochondrial genome (i.e., to be “petite positive”) (20). These observations raise the possibility that selective pressures related to carbon metabolism, and in particular transitions between fermentation and respiration, may have driven the retention of metabolic isozymes.

We wanted to determine whether isozyme pairs tend to act together as a functional unit, or whether, alternatively, each isozyme has a discrete role. If the former, then we would expect a strong tendency for isozymes to be co-expressed whereas otherwise we would expect anti-correlation or no correlation in expression otherwise. To address this question we assembled a large compendium of gene expression data consisting of more than 400 datasets (each comprising at least 6 arrays), and calculated the correlation of each isozyme gene pair’s expression within each dataset.

Unlike previous studies (see Note S1), we focused on statistically significant anti-correlation of gene expression within single datasets. We expect negative correlation of isozyme expression to be observed during experiments that capture the transition between environments where one vs. another isozyme is preferred. Further, when gene transcripts are measured by microarray, cross-hybridization can occur for highly homologous genes (24, 25), leading to artifactual positive correlation. Given that many isozymes in yeast have a large degree of homology, focusing on negative correlation mitigates this technical bias.

In our compendium, we found that overall, isozymes appeared to be anticorrelated less often than random gene pairs (Bonferroni-Holm-corrected Wilcox test *p* = 0.031), and more often than members of the same protein complex (p = 9.9×10^−5^) but did not differ significantly from other genes within the same metabolic pathway (p = 0.83; Figure 1a). When the correlation of isozyme pairs over the entire expression compendium was visualized, it became clear that this intermediate level of anticorrelation could be explained by the existence of two distinct clusters of isozymes: a minority of isozyme pairs appeared to be highly correlated across most of the compendium, while a majority showed strong anticorrelation under a subset of conditions (Figure 1b). Based on how often (i.e. in how many experiments) an isozyme pair was observed to show anticorrelated expression (q-value ≤ 0.1), we used logistic regression (see Methods) to classify the pair as either more like genes from the same protein complex (consistent with a role in dosage) or more like a pair of randomly-selected genes (suggesting independent roles for the individual isozymes). We found that 19 isozyme pairs resembled random pairs ≥10x more closely than they resembled pairs drawn from the same protein complexes; at the same threshold, 13 pairs more closely resembled members of the same protein complex (Supplemental Figure S2).

**Figure 1.**
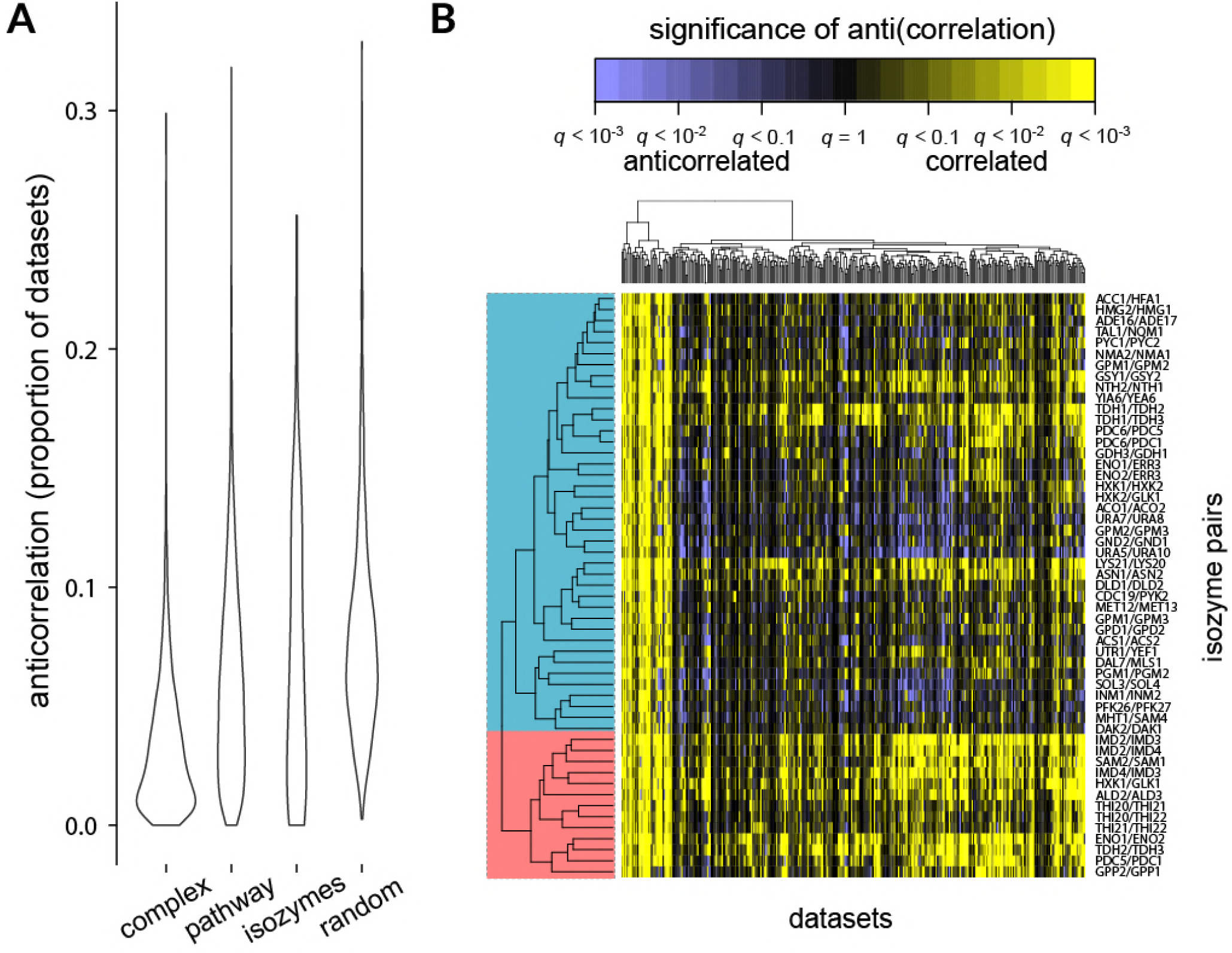
Many isozyme pairs are differentially expressed. a) Box plots of anticorrelation among isozyme pairs, compared with i) members of the same protein complex, ii) members of the same metabolic pathway, and iii) random gene pairs. Isozyme pairs are more likely to show differential expression than genes in the same complex (Bonferroni-Holm-corrected 2-sided Wilcox test *p*-value 9.9×10^−5^), but less likely than random genes (Holm-corrected p = 0.031). b) Isozyme pairs separate into two broad categories, depending on how often they are anticorrelated. The matrix displayed shows the correlation (yellow) or anticorrelation (blue) of isozyme pairs (rows) over every dataset (columns) in the compendium. Intensity corresponds to significance of the (anti)correlation (*q*-value). Hierarchical clustering using uncentered Pearson’s correlation reveals two main clusters of isozyme pairs: a minority are strongly correlated over most of the compendium, while a majority show condition-dependent anticorrelation.

As described in the Introduction, one explanation for isozyme retention is gene dosage: that is, having multiple copies of an enzyme may enable increased total enzyme expression (26). If isozymes were retained strictly for the purpose of increased dosage, we would not expect them to be differentially expressed. The prominence of anti-correlated pairs therefore demonstrates that dosage alone does not explain the continued retention of the majority of retained isozyme pairs, contrary to some previous assertions (2) but in accord with Ihmels et al. (27). Additionally, there was little overlap between co-expressed isozymes and those isozymes catalyzing high-flux reactions, as defined in a previous study (3), further arguing against a predominant role for dosage: only the GAPDH (*TDH1-3*) and hexokinase (*HXK1/GLK1*) enzymes appeared in both lists. Indeed, it appears that a majority of isozyme pairs are strongly anticorrelated in a condition-dependent manner, suggesting a role for these pairs in adaptation to different environments.

### A set of 21 isozyme pairs shows strong differential expression with changing carbon availability

Visualizing the anti-correlation of isozyme pairs also revealed that many were differentially expressed in the same datasets. This suggested that specific experimental conditions may be particularly relevant to explaining isozyme retention. We therefore wanted to identify the specific experimental perturbations leading to isozyme expression anticorrelation. Building on related work (5, 19, 27) (see SI Discussion), we first simply sorted the transcriptional datasets (each containing several individual arrays; for example, a heat shock time course would be one “dataset” (28)) by the number of isozyme pairs in each that were anticorrelated. Datasets with the most differential expression of isozyme pairs included many experiments related to the carbon source (Table S2). An alternative analysis by partitioning around medoids (PAM) clustering of the differential expression matrix revealed similar results (Supplemental Figure S3).

To test the association between carbon source perturbations and differential isozyme regulation more systematically, we performed dimensionality reduction of the datasets by grouping them into clusters of experiments in which the same genes showed the strongest expression changes. To accomplish this, we took the variance of each gene within each dataset and then clustered these variance vectors using a consensus *k*-means clustering, with the number of clusters determined by AIC (29) (see Methods). This method was effective at grouping together datasets reflecting similar experimental perturbations. For example, one cluster of datasets included diauxic shift time courses (30–32), carbon starvation time courses (33), a panel of mutants with and without glucose (34), and a 15-day wine fermentation (35). We then used this clustering to ask two questions: first, whether isozyme pairs were anticorrelated in a particular cluster, and second, whether they were more strongly anticorrelated within that cluster than in other datasets (see Figure 2a and Methods).

**Figure 2.**
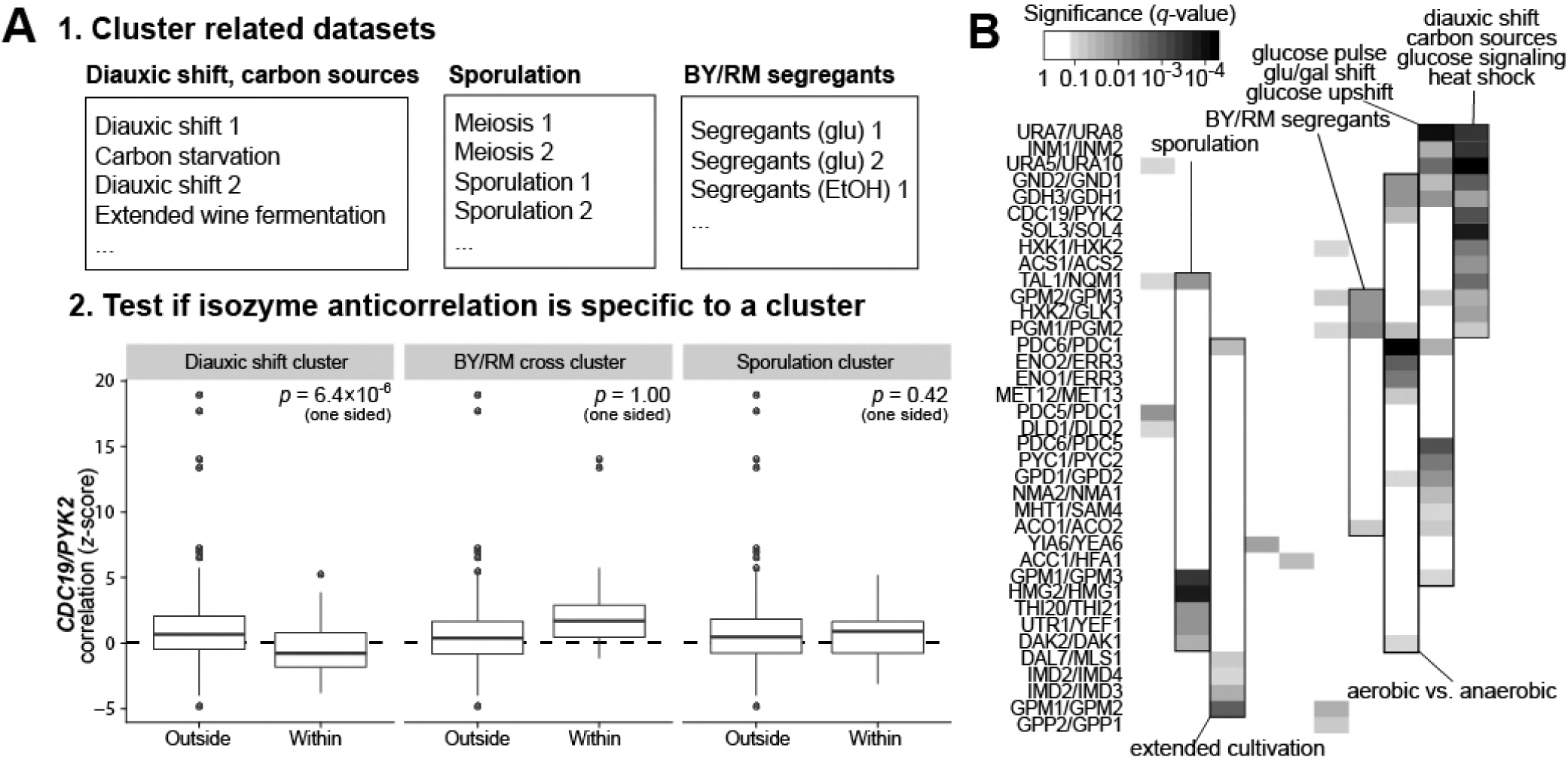
Sixteen isozyme pairs are associated with the metabolism of alternative carbon sources. a) Outline of method for association of isozyme pairs with particular dataset clusters. First, datasets are grouped into clusters of related experimental conditions (see Methods). Three of the resulting clusters are shown, with representative datasets. Next, for each dataset cluster and each isozyme pair, we test whether that pair is anticorrelated within that cluster, and if so, whether it is significantly more anticorrelated within that cluster vs. in other datasets. We show as an example *CDC19/PYK2*, which passes these criteria only within the first cluster of datasets (related to diauxic shift). This suggests the pair *CDC19*/*PYK2* is associated with the respirofermentative transition. b) A set of 16 isozyme pairs is specifically differentially expressed in a cluster of datasets having to do with metabolism of alternative carbon sources. Separately, 5 other pairs appear to be associated with sporulation and meiosis. A filled cell indicates that significant anticorrelation of a given isozyme pair was observed within a given dataset cluster, with the intensity of the cell corresponding to the *q*-value (the false discovery rate analog of a *p*-value).

Indeed, we found that a core set of 13 isozyme pairs tended to be particularly strongly anticorrelated in a cluster of conditions having to do with diauxic shift/glucose limitation, and a partially-overlapping set of 14 pairs was strongly anticorrelated in a cluster of datasets containing several glucose pulse/upshift experiments; these pairs number 21 in total (Figure 2b). We also found that, for instance, 6 isozyme pairs were specifically associated with meiosis and sporulation, and 9 pairs with aerobic vs. anaerobic growth. These findings highlight the ability of this method, when applied to a large expression compendium, to associate sub-groups of anticorrelated isozymes not only with stress in general, but also to more specific environmental stressors.

Examining the original expression data from diauxic shift and glucose removal experiments revealed a clear visual pattern of anticorrelation (Figure 3a), which was conserved across different yeast strains (Supplemental Figure S4) (36) and even across species as shown using data from the most diverged *Saccharomyces sensu stricto* yeast, *Saccharomyces bayanus* (now called *Saccharomyces uvarum)* (Figure 3b) (37). Taken together, these results suggest that a core set of central carbon metabolic isozymes may be involved in adaptation to non-fermentable carbon sources.

**Figure 3.**
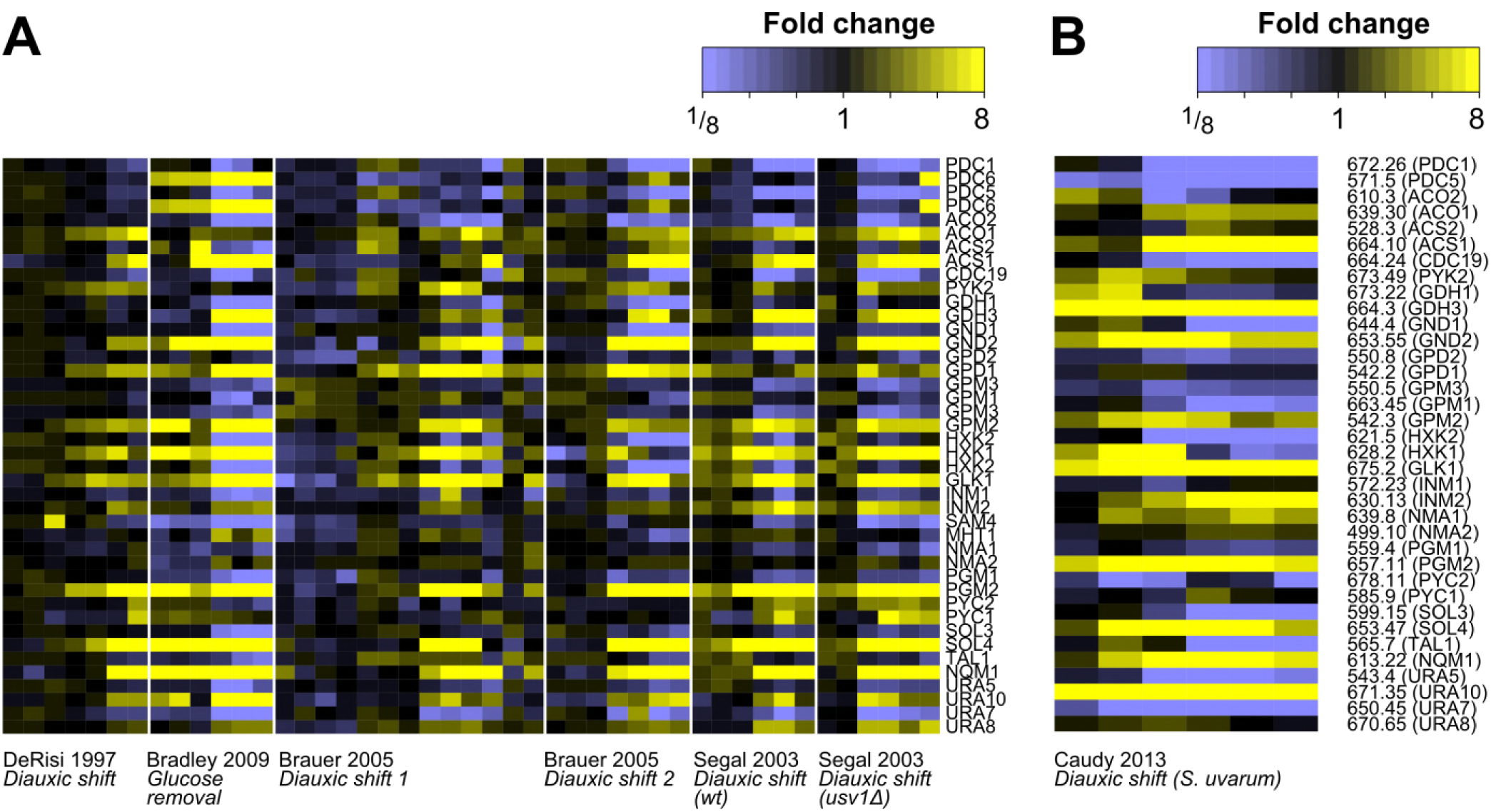
Anticorrelation of 16 isozyme pairs in response to glucose availability. a) Gene expression profiles of isozymes that are associated with the transition from using glucose to using alternative carbon sources. Array data from several diauxic shift and carbon removal experiments were collected, showing induction of one member of the pair (yellow) and repression of the other (blue) across the diauxic shift. Intensity corresponds to fold change. Genes are grouped into isozyme pairs. b) Gene expression signature of isozymes is conserved over evolutionary time. Isozymes were mapped to their syntenic orthologs in *Saccharomyces uvarum.* The expression of these orthologs in *S. uvarum* during the diauxic shift (86) shows the same overall pattern as the original isozymes in *Saccharomyces cerevisiae* (compare panel *a*).

We next sought to assign function to “minor” metabolic enzymes, using the aconitase isozyme *ACO2* as an example of an isozyme that is selectively expressed when glucose is available, and the pyruvate kinase isozyme *PYK2* of an example of the converse, an isozyme selectively expressed in the absence of glucose. Additionally, both *ACO2* and *PYK2* have isozyme paralogs (*ACO1* and *CDC19*) with profound deletion phenotypes, but have subtle (*ACO2*) or no (*PYK2*) recorded deletion phenotypes themselves.

### Aconitase 2 is required for efficient glycolytic respiration

Aconitases are iron-sulfur proteins that catalyze the second step of the TCA cycle, taking citrate to its isomer, isocitrate, via aconitate. This reaction does not require redox or nucleotide cofactors, nor is it at a branch point in metabolism; however, it is required for α-ketoglutarate synthesis and TCA cycle turning.

Yeast has two aconitase isozymes, *ACO1* and *ACO2*, both of which are mitochondrial; deletions of *ACO1* and *ACO2* are synthetically lethal (13). From the above analysis of microarray data, we noticed that *ACO1* is repressed by glucose and expressed on glucose removal, while *ACO2* has the opposite transcriptional pattern. *ACO1* is the “major” isozyme, and its deletion has been shown to be severely defective on respiratory carbon sources, such as glycerol, ethanol, and lactate (38). Its expression in the absence of glucose is consistent with the activation of TCA turning. Given that yeast prefer to ferment in the presence of glucose, the function of the *ACO2* isozyme was unclear, although a high-throughput competitive fitness screen had reported that an *aco2Δ* strain had a growth defect in minimal medium with glucose (14).

We began our experimental studies with an *aco2Δ* mutant strain by studying its growth in glucose minimal medium. We observed no growth defect during exponential phase in glucose minimal medium, indicating that residual expression of *ACO1* is sufficient to support synthesis of α-ketoglutarate and associated amino acid products (e.g., glutamate, glutamine, lysine). Growth of the *aco2Δ* deletion strain, however, saturated earlier than wild-type in glucose minimal medium (Figure 4b, inset). This suggests that *ACO2* plays an increasingly important role as glucose becomes limiting.

**Figure 4.**
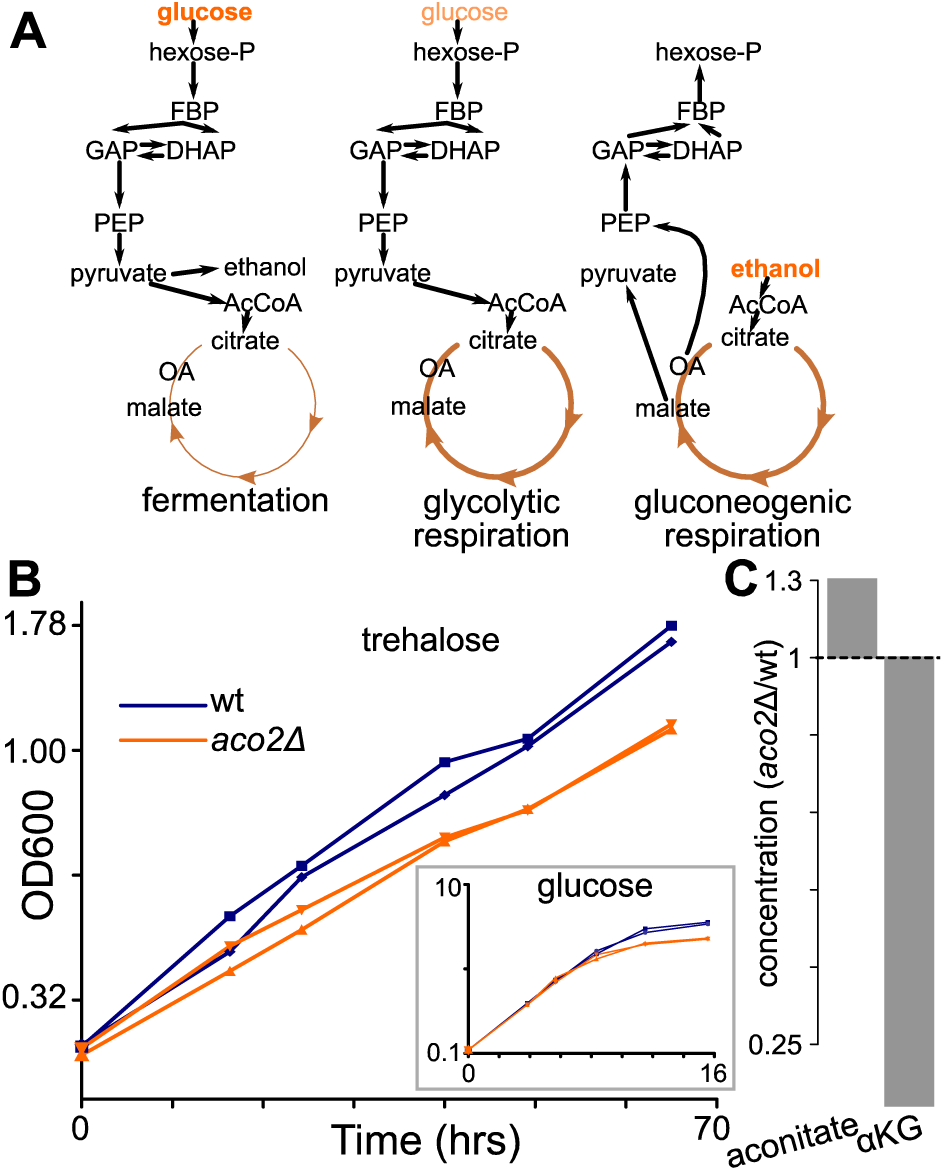
Deletion of the minor aconitase isozyme *aco2* results in a selective growth defect on trehalose, indicating impaired glycolytic respiration. a) Schematic of metabolism across the diauxic shift. In the presence of high levels of glucose (left), *S. cerevisiae* prefers to ferment glucose to ethanol. As glucose becomes limiting (center), *S. cerevisiae* continues to use glucose but converts it into acetyl-CoA and eventually CO_2_, in so doing driving TCA cycle turning and oxidative phosphorylation. We term this state glycolytic respiration. Finally, when glucose is exhausted, *S. cerevisiae* uses ethanol to make acetyl-CoA, as well as sugar phosphates through gluconeogenesis. We refer to this state as gluconeogenic respiration. b) Growth of wild-type and *aco2Δ* strains on minimal medium with trehalose, which is digested extracellularly to provide a steady but limiting amount of glucose, reveals a growth defect for *aco2Δ* during gluconeogenic respiration. In contrast, when grown on glucose (inset), the *aco2* deletion mutant has no growth defect in log-phase and only begins to show a growth defect when glucose becomes limiting. Data are biological duplicates. c) During steady-state growth on limiting glucose, aconitate is slightly elevated (137% of wild-type) and α-ketoglutarate decreased (56% of wild-type) in the *aco2Δ* mutant compared to wild-type. Bar plots represent averages of four technical replicates (repeated sampling from one chemostat per strain).

On limiting glucose, wild-type *S. cerevisiae* continues to perform glycolysis but, instead of fermenting the resulting pyruvate to ethanol, activates respiration to make more ATP. We refer to this state as “glycolytic respiration,” as distinguished from “gluconeogenic respiration,” in which cells respire using 2- and 3-carbon substrates like ethanol, glycerol, or acetate (Figure 4a). We hypothesized that the function of the *ACO2* isozyme is to support glycolytic respiration. To test this hypothesis, we grew the *aco2*Δ strain on minimal medium with trehalose as the carbon source. Trehalose, a glucose-glucose disaccharide, is cleaved extracellularly by *S. cerevisiae*; this produces glucose at a slow rate (39), inducing sustained glycolytic respiration. On trehalose, the *aco2*Δ deletion had a fitness disadvantage of 25% (Figure 4b), confirming that this aconitase isozyme supports glycolytic respiration. Furthermore, we observed no defect of the *aco2*Δ deletion when grown on minimal media with gluconeogenic carbon sources (Supplemental Figure S5), indicating that the metabolic role of *aco2*Δ is specific to glycolytic and not gluconeogenic respiration.

We also profiled the metabolome of *aco2*Δ and compared it with wild-type, using chemostat culture to maintain steady state growth on limiting glucose. Consistent with lowered aconitase activity, aconitate levels were somewhat elevated and α-ketoglutarate depleted (Figure 4c). We also observed increases in the levels of compounds in the *de novo* NAD^+^ biosynthesis pathway from tryptophan, such as kynurenic acid (Supplemental Figure S6). This connection to NAD^+^ biosynthesis aligns with previous observations that a deletion of *bna1* (a key NAD^+^ biosynthetic gene) is synthetically sick with *aco2*Δ (40). Further work is required to identify the molecular mechanism underlying this phenotype.

### Pyruvate kinase 2 is required for efficient growth on three-carbon substrates

We next studied growth of the pyruvate kinase isozyme encoded by the *PYK2* gene, an example of an isozyme that is selectively expressed in the absence of glucose. Pyruvate kinase catalyzes the last step of glycolysis, taking phosphoenolpyruvate (PEP) to pyruvate and producing ATP from ADP. This step of glycolysis is highly regulated from yeast (41) to humans (42). The “major” yeast isozyme is known as *CDC19* (where “CDC” is from “cell division cycle”: a *cdc19* deletion causes arrest at the G1/S transition). It is expressed in the presence of glucose and its activity requires high cytosolic fructose-1,6-bisphosphate levels, which are produced when glucose is abundant. The *PYK2* isozyme lacks such regulation by fructose-1,6-bisphosphate. Deletion of *CDC19* is lethal on glucose, but deletion of *PYK2* has no known phenotype on either glucose or ethanol (43). Because it does not require activation by FBP, it has been suggested that *PYK2* may contribute to fitness specifically when glucose is limiting (44); however, we found that the *pyk2*Δ deletion exhibits no growth defect on trehalose, contradicting this hypothesis (Figure 5a). While ethanol-fed cells must also make pyruvate, they appear to do so primarily from the TCA cycle via malic enzyme (*MAE1*) (43), rendering pyruvate kinase unimportant.

**Figure 5.**
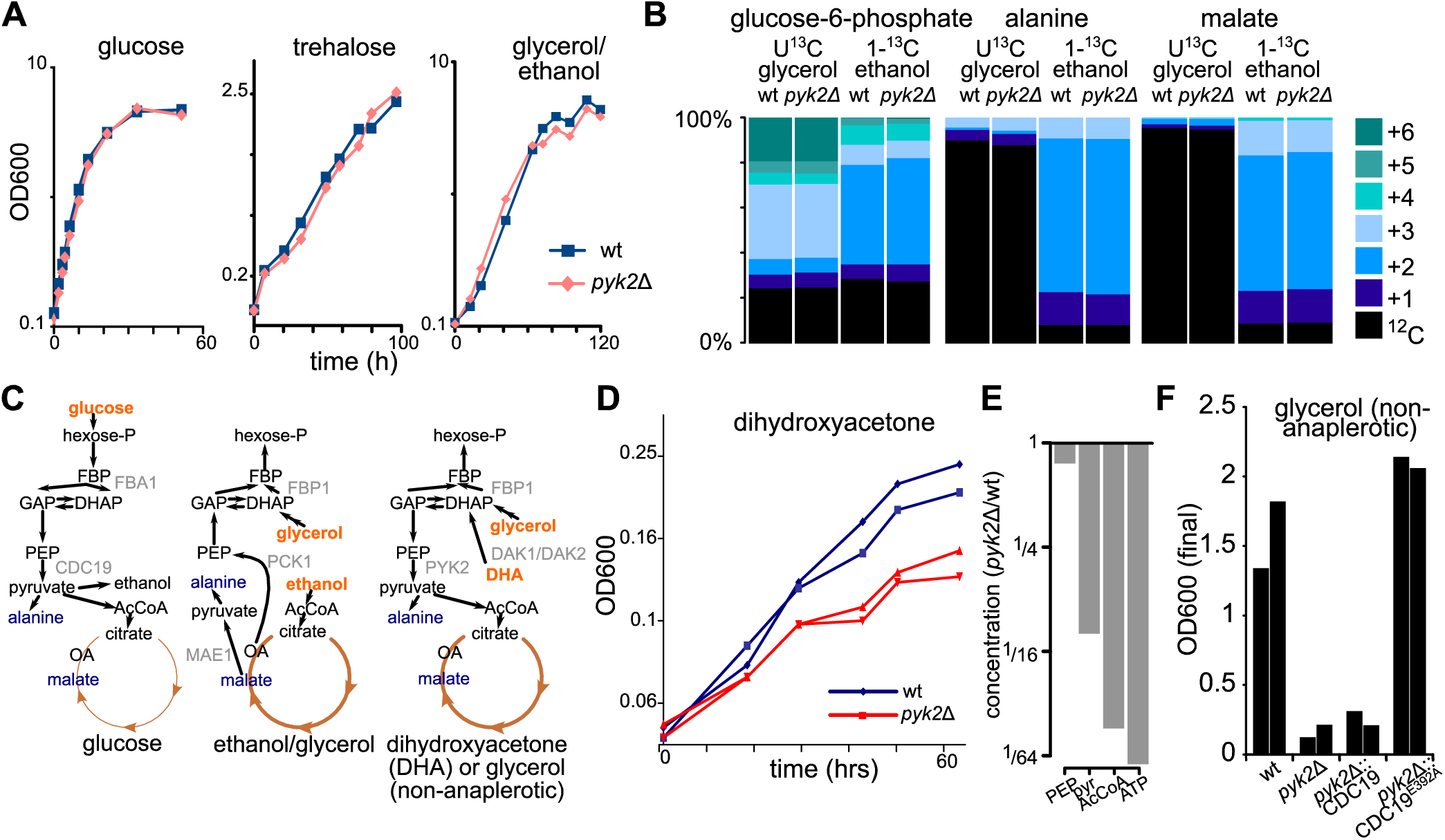
Deletion of the minor pyruvate kinase isozyme *PYK2* results in a selective growth defect on dihydroxyacetone, a three carbon sugar. a) Growth of wild-type and *pyk2Δ* strains on glucose, trehalose, and glycerol/ethanol minimal media, revealing no defect for *PYK2* deletion. b) ^13^C-labeling shows that, for wild-type yeast growing on glycerol/ethanol minimal medium, glucose-6-phosphate labels from both glycerol and ethanol, but alanine (a proxy for pyruvate) and malate are labeled exclusively from ethanol. Thus, glycerol is not used to make pyruvate. The *PYK2* deletion does not affect these labeling patterns. Labeling patterns are averages of biological duplicates. c) Schematic of glycolysis and TCA cycle comparing the metabolism of glucose, glycerol/ethanol medium, and dihydroxyacetone. d) Growth of wild-type and *pyk2Δ* strains on dihydroxyacetone minimal medium reveals a growth defect for *pyk2Δ*. Data are biological duplicates. e) At steady state on limiting dihydroxyacetone, phosphoenolpyruvate changes only slightly in concentration (85% of wild-type), while pyruvate (down 3.4-fold), acetyl-CoA (down 12-fold), and ATP (down 16-fold) are substantially decreased. Bar plots represent averages of four technical replicates (repeated sampling from one chemostat per strain). f) Growth on synthetic glycerol (glycerol/CSM-Arg-Asp) medium is normal in wild-type and when *pyk2Δ* is rescued with the FBP-insensitive mutant *CDC19E392A*, but near-abolished for *pyk2Δ* and *pyk2Δ* rescued with wild-type *CDC19*. Bars are final OD600 of biological duplicates.

*S. cerevisiae* grows better on a mixture of glycerol and ethanol than ethanol alone. While this glycerol could potentially be used to make pyruvate via PEP, the *pyk2*Δ deletion displayed no growth phenotype on glycerol/ethanol (Figure 5a). This raised the possibility that in cells fed glycerol-ethanol, like cells fed ethanol alone, pyruvate is made via *MAE1*. We confirmed this via experiments with ^13^C-glycerol and LC-MS: upon feeding uniformly ^13^C-labeled glycerol and unlabeled ethanol, while glucose-6-phosphate labeled from glycerol as expected, alanine (which is made by transamination of pyruvate) remained primarily unlabeled (Figure 6b). Conversely, feeding the cells [1-^13^C]-ethanol led to labeling in both malate and alanine. This indicates a lack of reliance on pyruvate kinase in glycerol-ethanol fed cells. Indeed, the *pyk2*Δ deletion showed no difference from wild-type in labeling patterns (Figure 5b).

In light of the above results, we hypothesized that *PYK2* would be required on carbon sources that (i) result in insufficient fructose-1,6-bisphosphate levels to activate *CDC19* and (ii) require pyruvate kinase activity to make pyruvate. We further reasoned that 3-carbon substrates would meet the above requirements. While glycerol is a common 3-carbon substrate, *Saccharomyces cerevisiae* cannot grow on glycerol minimal medium without amino acids, presumably because of inability to maintain cytosolic redox balance (glycerol is more reduced than glucose) (45). We therefore searched for another 3-carbon substrate that could sustain growth as the sole carbon source. *S. cerevisiae* has two dihydroxyacetone kinases, *DAK1* and *DAK2*, that enable slow but sustained growth on the triose dihydroxyacetone (DHA) (46). DHA enters metabolism through glycolysis/gluconeogenesis, as opposed to through the TCA cycle, so during growth on DHA, pyruvate should be made using pyruvate kinase as opposed to malic enzyme. Since *CDC19* is turned off in the absence of glucose (both transcriptionally and allosterically), the majority of the flux through pyruvate kinase should be catalyzed by *PYK2* (Figure 5c).

Indeed, we observed that deletion of *pyk2* inhibited growth on DHA (Figure 5d). Further, during continuous culture on dihydroxyacetone, while phosphoenolpyruvate (PEP) levels remained close to wild-type, the *pyk2*Δ deletion was depleted in pyruvate kinase’s products: pyruvate and ATP. The downstream pyruvate product, acetyl-CoA, was also depleted. This pattern of metabolite levels is consistent with impaired pyruvate kinase activity in this mutant (Figure 5e; see also Supplemental Figure S6).

Further, while lab yeast do not grow on a strict minimal medium with glycerol, amino acid supplementation can restore growth. For example, the “complete supplement mixture” (CSM), containing selected amino acids and nucleobases, is sufficient to permit growth on glycerol (45). In *Saccharomyces cerevisiae*, amino acid degradation does not always yield carbon skeletons that can enter central carbon metabolism; instead, the carbon skeletons of many amino acids are either used only in amino acid biosynthesis, or are discarded in the form of fusel alcohols via the Ehrlich pathway, which has been speculated to play an important role in maintaining redox balance (47). The only amino acids in CSM known to be catabolized to central carbon intermediates are L-aspartate and L-arginine. To allow growth on glycerol without the confounding influence of other potential carbon sources, we therefore constructed a synthetic glycerol medium supplemented with a version of CSM lacking these amino acids (CSM-Arg-Asp).

When grown on glycerol with CSM-Arg-Asp, we observe that the *pyk2*Δ deletion strain is severely impaired relative to wild-type (Figure 5f). In contrast, previous studies have found no defect of *pyk2*Δ during growth on synthetic complete media containing all amino acids. Together, these results indicate a previously unappreciated role for the pyruvate kinase isozyme *PYK2* in allowing metabolism of three-carbon substrates.

Finally, we wanted to definitively test our hypothesis about the mechanism by which *PYK2* permits growth on three-carbon substrates: that is, that *PYK2* enables growth on glycerol and dihydroxyacetone because unlike *CDC19*, it does not require allosteric activation by FBP, which is depleted in the absence of glucose. Since *CDC19* is repressed under non-fermentative conditions, however, it could also be possible that either pyruvate kinase allows growth on 3-carbon conditions and that the relevant difference between *PYK2* and *CDC19* is regulation at the promoter level.

To distinguish between these two possibilities, we performed two rescue experiments in which, using the *delitto perfetto* method (48), we kept the promoter of *PYK2* completely intact, but replaced the *PYK2* coding sequence with either the wild-type *CDC19* or the E392A point-mutant of *CDC19*, which allows *CDC19* activity regardless of FBP concentrations (49, 50). Rescue with wild-type *CDC19* did not improve growth on glycerol with CSM-Arg-Asp; rescue with the E392A allele, in sharp contrast, restored growth completely (Figure 6f). Similar results, with a partial rescue by the wild-type *CDC19* allele and a complete rescue by the E392A allele, were also observed for dihydroxyacetone (Supplemental Figure S7). These results support our hypothesis that the *PYK2* gene has been retained in *Saccharomyces cerevisiae* because of an “escape from adaptive conflict” (51–53): the presence of *PYK2* allows the cell to control *CDC19* activity through allosteric activation, which has previously been shown to be important for adapting to short-term glucose removal (54), while also resolving the incompatibility between this regulation and growth on three-carbon substrates.

## Discussion

Through computational analysis of a large compendium of expression data, we found that a set of co-localized metabolic isozymes is differentially expressed in response to glucose availability, and that this differential expression is conserved over evolutionary time. We then experimentally found condition-specific contributions to fitness for two of these isozymes, *ACO2* and *PYK2.* In the case of *ACO2*, the deletion shows only a subtle defect on glucose but a defect of 25% on trehalose (i.e. limiting glucose). In the case of *PYK2*, the deletion shows no defect on glucose, ethanol, or glycerol-ethanol, but a defect of 42% on dihydroxyacetone, and near-complete growth inhibition when glycerol was the sole carbon source. Also, another study confirmed that the acetyl-CoA synthase isozymes *ACS1* and *ACS2*, identified as part of the same cluster of differentially-expressed isozymes returned using our method, display opposite phenotypes on fermentable and non-fermentable carbon sources (55), further supporting the validity of this approach.

The conclusion that even so-called “minor” isozymes make important contributions to fitness that cannot be easily buffered is in line with other recent genome-scale analyses. One recent study first predicted fitness costs for gene deletions using a flux balance model, then calculated the evolutionary rates of these genes using sequence analysis, and finally asked whether genes with larger predicted deletion phenotypes evolved more slowly. The authors demonstrated this expected relationship only appeared when isozymes were assumed to be non-redundant, as opposed to individually dispensable (56). Furthermore, a study analyzing experimentally determined deletion phenotypes concluded that even closely-related, non-essential duplicates actually made distinct, condition-specific contributions to fitness, with effect sizes that are likely large enough for purifying selection to have retained them both (57). Finally, another study integrating experimental data with several bioinformatic estimates of functional divergence concluded that hardly any paralogous pairs are truly functionally redundant (58). Together, these lines of inquiry reinforce the conclusion that failure to find fitness differences in standard media does not indicate dispensability: the differences in fitness that led to retention of a gene may often be detected only under very specific growth conditions. As in the examples we provided here, careful analysis of the relevant biochemical pathways may often be required to infer the appropriate environments in which the differences in fitness can be made manifest.

Overall, out of 77 isozyme pairs in yeast, our bioinformatic analyses suggest that 24 are differentiated primarily by compartment, 21 by gene expression on carbon sources, and 16 by gene expression in other conditions. This leaves 16 whose expression is strongly positively correlated, suggesting a potential primary role for gene dosage effects. However, few of these 16 pairs catalyze high-flux reactions according to Kuepfer et al. (3), suggesting that even in these cases, dosage may not be the primary explanation. Furthermore, some metabolic isozymes that do not show gene expression differences in our analysis have been reported to be differentially regulated at the level of protein concentration. For example, *ENO1* and *ENO2*, the cytosolic enolases, display opposite changes in abundance due to glucose availability when assayed by chromatography followed by activity assay (59). This is consistent with the observation that deletion of *eno2*, but not *eno1*, causes abnormal cell cycle progression during standard growth (60). However, *ENO1* and *ENO2* are highly correlated across our expression compendium. This may be either because their mRNAs are hard to distinguish by microarray, or because their primary regulation is at the level of translation, post-translational modification, or protein stability. Greater availability of RNA sequencing and quantitative proteomics data will therefore be valuable for this type of analysis in the future.

For isozymes that are differentiated by condition-specific expression, dosage may still have played an important role in their initial evolutionary selection. For example, Conant and Wolfe argue that loss of duplicates outside of glycolysis may have led to higher flux through glycolysis and thus a competitive fitness advantage shortly following the whole genome duplication in yeast (26). In this model, further specialization of isozymes occurred shortly following the whole-genome duplication. Such further specialization likely provided an evolutionary benefit through escape from adaptive conflict: gene duplication allows the expression level and/or enzymatic activity of a protein to be tailored to two different conditions, where a single protein would have to “split the difference” and would thus be imperfectly adapted to both (52).

What kinds of adaptive conflict might drive isozyme differentiation? One conflict is between affinity and speed (K_m_ vs. k_cat_), as exemplified by the hexose transporters. Another kind of conflict arises from differing allosteric regulatory requirements, as exemplified by the pyruvate kinases. Allosteric regulation of pyruvate kinase by fructose-1,6-bisphosphate (FBP) is conserved from bacteria (61) to human (62). Recently, it has been recognized that ultrasensitive (i.e., cooperative) activation by FBP enables pyruvate kinase activity to turn “on” and “off” in a switch-like manner in response to glucose availability (54). In bacteria, similar allosteric activation has also been observed for not only pyruvate kinase but also the other main PEP-consuming enzyme, PEP carboxykinase (50). Such switch-like regulation facilitates growth in oscillating glucose environments and prevents futile cycling in gluconeogenic ones. It is problematic, however, for substrates that enter metabolism via lower glycolysis. They must rely on simultaneous “downward” flux through pyruvate kinase, on the one hand, and “upward” flux through fructose-bisphosphate aldolase to produce 6-carbon sugars, on the other. “Downward” flux requires high FBP to activate pyruvate kinase, while “upward” flux requires low FBP to render net FBP formation by aldolase thermodynamically favorable. Here, we show that the solution involves expression of *PYK2,* whose activity does not depend on high FBP levels.

It is notable that even though the differential allosteric regulation of *CDC19* and *PYK2* has been known for 20 years, and despite extensive interest in pyruvate kinase isozymes due to the strong association of the mammalian pyruvate kinase M2 isozyme with cancer (63), no functional role for *PYK2* had been previously identified. Indeed, a recent competitive fitness study assaying more than 400 growth conditions revealed a growth phenotype on at least one condition for 97% of yeast genes, but did not find any fitness defect for the *pyk2*Δ deletion (16).

Another study (64) expressed both *CDC19* and *PYK2* from non-native promoters to tune pyruvate kinase function experimentally, concluding that lower pyruvate kinase activity was accompanied by an increase in oxidative metabolism and oxidative stress resistance. This is in line with previous reports that slower growth rates, as the authors show occurs with pyruvate kinase downregulation, induce both respiration and stress-protective machinery in yeast (18, 65) which includes intracellular glutathione, an important antioxidant (66). However, this work does not give a specific functional role for *PYK2*, and does not provide experimental evidence explaining the retention of *PYK2* in wild-type yeast. Here, we demonstrate that the Pyk2p protein product specifically, and not Cdc19p, is important for efficient growth on three-carbon substrates.

Despite decades of research on yeast physiology, dihydroxyacetone was only identified as a sole carbon source for *S. cerevisiae* in 2003 (46). Further, even though glycerol is a commonly-used non-fermentable carbon source in yeast biology, it has almost always been used in rich or extensively supplemented media, which contain other potential carbon sources (45). A full understanding of the role of yeast isozymes in central metabolism will likely go hand-in-hand with a more complete understanding of potential modes of carbon metabolism. More generally, many functions of yeast metabolic genes and enzymes may only become manifest when growth and viability in a wider variety of environments are studied. Because prolonged propagation on glucose may have led to the loss of certain metabolic capabilities (e.g., xylose catabolism) in lab yeast strains (67), study of natural isolates may also be important.

An important dichotomy in our findings is between the conditions that control isozyme expression, and those in which isozymes are functionally important. A similar duality between the genes induced under a particular condition and the genes necessary for growth in that condition has been previously observed (68–70). Here, we find that the expression of a large subset of isozymes is controlled by glucose availability, and indeed experimentally confirm that two “minor” isozymes play important roles when glucose is low or absent. Yet this broad characterization in terms of gene expression belies much more complex and specific functional roles for these isozymes. For example, *ACO2* is expressed most in the presence of glucose, yet *ACO2* is functionally important only when glucose is scarce: this indicates that its expression in high glucose actually reflects preparation for future times when glucose is limiting. Similarly, the absence of glucose induces expression of *PYK2*, yet *PYK2* is useful only in the case of carbon sources that feed into lower glycolysis, bypassing the metabolite (and key allosteric regulator) fructose-1,6-bisphosphate. Thus, a set of isozymes render yeast central carbon metabolism more flexible, allowing a small number of fundamental transcriptional states to produce optimal enzyme activities across a broad range of potential environmental conditions.

## Methods

### Identifying metabolic isozymes of the same compartment

Our criteria for identifying isozymes relevant to our study were as follows: (i) The proteins had to perform the same reaction: we drew our initial list of protein pairs meeting these criteria from the reconstructed metabolic model of *Saccharomyces cerevisiae* iLL672 (3); (ii) The proteins needed to have the same small molecules as products and reactants, as detailed in the Yeast Pathway Database (23); we therefore excluded, for example, the protein mannosyl-O-transferases *PMT1-6*, since these enzymes have different proteins as substrates; (iii) We included only isozymes annotated as preferring the same co-factors (e.g. NADP^+^ vs. NAD^+^) in the Saccharomyces Genome Database (71); (iv) We only considered isozymes whose products and reactants were in the same subcellular compartment (i.e., excluding transporters); (v) We excluded isozymes that were annotated to different compartments (e.g. mitochondria vs. cytosol) (72) (24 pairs). Our final list was comprised of 53 isozyme pairs and 85 genes in total, some having more than one partner (e.g., the three glyceraldehyde-3-phosphate dehydrogenases *TDH1*, *TDH2*, and *TDH3*).

To test whether pathways were enriched for isozymes, pathways were drawn from the Yeast Pathway database (23) and then combined into the categories shown in Supplemental Figure S1. (When testing “central carbon metabolism,” we included all reactions from “glycolysis, gluconeogenesis, fermentation”, “pentose phosphate pathway”, and “TCA cycle.”) The proportion of reactions catalyzed by isozymes in each category was calculated, and the two-tailed Fisher’s exact test was used to establish significance (Table 1).

### Assessing anticorrelation of metabolic isozymes

#### Processing of gene expression data

Microarray data from the Gene Expression Omnibus (GEO) (73) and SPELL (28) were downloaded, processed, and divided into single-experiment datasets as in Note S2 (Supplemental Methods). We then constructed an m × n binary matrix of anticorrelation B such that each entry b_m, n_ was 1 if gene pair m was anticorrelated at a significance threshold of q < 0.1 in dataset n. The matrix B was sorted by columns from most to least anticorrelation (Σ_m_b_m, n_); the top 10 datasets with the most anticorrelated pairs are listed in Supplemental Table S2. To determine whether isozyme pairs separated naturally into multiple groups, B was also clustered using partitioning around medoids (PAM) with k = 3 clusters, yielding two coherent clusters (Note S2).

#### Comparison of isozymes with other types of proteins

We compared differential expression of isozymes with other pairs of genes. Lists of genes in protein complexes came from a high-throughput pulldown/mass-spectrometry assay; only “core” complexes (i.e., sets of proteins that co-purified most often) were used (74). We then computed the proportion of arrays in which a given gene pair was differentially expressed, *p*_*m*_ = (Σ_*n*_*b*_*m*, *n*_)/(*n*). Here, as above, *b*_*m*, *n*_ = 1 if the correlation *r*_*x*, *y*, *d*_ was significantly less than 0 at a q-value of 0.1 (75), and was set to 0 otherwise; row *m* corresponds to gene pair (*x*, *y*) and column *n* corresponds to dataset *d*. The distributions of *p*_*m*_ values were compared via the nonparametric two-tailed Kolmogorov-Smirnov test.

#### Testing differential expression within dataset clusters

To cluster datasets, per-gene standard deviations were computed for every dataset. Missing values in the resulting *m* × *n* matrix, with m as the number of genes and n as the number of datasets, were imputed using KNNimpute with 10 neighbors (76), discarding first genes and then datasets with more than 70% missing values. Standard deviations were logged after adding a constant equal to half the smallest standard deviation in a given dataset. Next, the matrix was first column-normalized (i.e. mean-subtracted and divided by standard deviation) and then row-normalized, to ensure that genes or datasets with larger dynamic ranges did not dominate the clustering. This is a related approach to that described by Tavazoie et al. (77).

To ensure robustness of the clustering, a consensus *k*-means clustering (29) was then performed for *k* ranging from 2 to 50. Briefly, in this consensus clustering, 125 subsamples of the original matrix were generated, sampling 80% of the rows and 80% of the columns. These subsamples were clustered via *k-*means, and the resulting clusterings were converted into a “consensus matrix” giving the proportion of subsamples in which two datasets clustered together; this consensus matrix was then hierarchically clustered and cut to give *k* groups. For each value of k, AIC was calculated (i.e., RSS(*k*) + 2*Mk*, where RSS is the residual sum of squares with *k* clusters and *M* is the length of each vector) (78). AIC was then minimized, yielding *k* = 16 clusters.

To test for specificity of differential expression within dataset clusters, for each isozyme pair (*i*_1_, *i*_2_) and cluster of datasets *C*, we stipulated two criteria. First, we required that the average normalized correlation of the pair (*i*_1_, *i*_2_) tended to be negative in datasets *d* within the cluster *C*, i.e.,

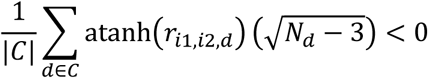

(Here, *r*_*i*1, *i*2_ is the Pearson correlation of isozymes *i*_1_ and *i*_2_, and *N*_*d*_ is the number of observations in dataset *d*. *atanh* is the hyperbolic arctangent function, used for the Fisher z-transform as described in 4.2.1.) Second, we tested whether the normalized correlation of the pair 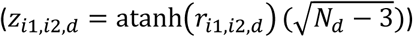 tended to be less within the cluster than without (i.e., *z*_*i*1, *i*2, (*d* ∈ *C*)_ < *z*_*i*1, *i*2, (*d*∉*C*)_), using a one-tailed rank sum test. *p* values for this test were corrected according to the *q*-value method of Storey (75), and a cutoff of q ≤ 0.1 was applied.

### Strain construction, media and growth conditions

#### Strains

Prototrophic *aco2* and *pyk2* deletion strains were provided by David Hess and Amy Caudy from their prototrophic deletion collection (79) The final prototrophic deletion strains had the genotype MAT*α*, *yfg*Δ::*KanMX*, *can1Δ::STE3pr-SpHIS5*, *his*3Δ0, *lyp*1Δ, where *yfg* represents either *aco2* or *pyk2*. For the rescue experiments, wild-type *CDC19* or *cdc19-*E392A were introduced into the native *PYK2* promoter using the *delitto perfetto* allele-replacement method (48).Starting with DBY12000, a *HAP1*+ derivative of FY4, *PYK2* was knocked out using the *pCORE* construct, which contains an antibiotic resistance cassette (KanMX) and a counterselectable marker (*URA3*): selection for resistance to geneticin yielded a *pyk2*Δ*::CORE* strain. *URA3* was then knocked out to allow for use of the counterselectable marker. *CDC19* (wt or E392A) was then amplified using primers with overhangs homologous to this construct. The resulting PCR products were then transformed into the *pyk2*Δ::*CORE* knockout and transformants were selected based on loss of the counterselectable marker (i.e., resistance to 5-FOA), yielding pyk2Δ::*CDC19* and pyk2Δ::*cdc19-*E392A strains in a *ura3*Δ background. Finally, these strains were mated to a strain with wild-type *URA3*; sporulation and dissection yielded fully prototrophic strain with genotypes MAT**a**, pyk2Δ::*CDC19*(wt/E392A).

#### Media recipes

Minimal media for batch cultures were prepared using 6.7g/L Yeast Nitrogen Base (Difco) and an appropriate carbon source. Final concentrations of carbon sources were 20g/L for glucose (YNB-glucose), 100g/L for trehalose as per Jules et al. (39) (YNB-trehalose), 20g/L each for glycerol/ethanol (YNB-glycerol/ethanol), and 9 g/L (100 mM) for dihydroxyacetone as per Boles et al., 1998 (YNB-DHA). Minimal media for chemostats were prepared according to the glucose-limited chemostat media (CM-glucose) recipe in Dunham et al. (80); for DHA-limited chemostats (CM-DHA), 8 mM DHA was substituted for 8 mM glucose. Synthetic glycerol medium (glycerol/CSM-Arg-Asp) was prepared using YNB without ammonium and 3% glycerol, plus the following nitrogen bases and L-amino acids: 10 mg/L adenine, 20 mg/L His, 50 mg/L Ile, 100 mg/L Leu, 50 mg/L Lys, 20 mg/L Met, 50 mg/L Phe, 100 mg/L Thr, 50 mg/L Trp, 50 mg/L Tyr, 20 mg/L uracil, and 140 mg/L Val.

#### Batch cultures

Wild-type (*ho*Δ) and isozyme deletions (*aco*2Δ, *pyk*2Δ) were struck out on YPD. For growth curves, a different colony was picked for each biological replication and placed into YNB-glucose and grown overnight. Overnight cultures were then set back in YNB-glucose for at least one doubling. For growth curves on glucose, the cultures were then set back such that the optical density at 600nm (OD600) was close to 0.1. For growth curves on YNB-trehalose and YNB-dihydroxyacetone, log-phase cultures grown in glucose were set back into media containing either trehalose or dihydroxyacetone and allowed to double at least twice in the new medium; cultures were then set back to an OD600 of 0.1 to start the growth curve. For the experiment measuring final density on glycerol/CSM-Arg-Asp, cultures were inoculated into SD, then washed 3x in YNB without ammonium, and set back to an initial OD600 of 0.05 in glycerol/CSM-Arg-Asp; final OD600 was measured after 16 days.

#### Chemostat cultures

Wild-type and isozyme deletions were struck out onto a YPD plate and then inoculated in CM-glucose (for glucose-limited chemostats) or a minimal medium with YNB, dihydroxyacetone, and 0.05% glucose (CM-DHA-glucose, for DHA-limited chemostats). For CM-DHA-glucose, overnights were allowed to grow an additional day. Overnights were then used to inoculate one chemostat per strain/medium combination, with media limited for either glucose or for dihydroxyacetone (see media recipes). Batch mode proceeded for one day for glucose-limited chemostats and three days for DHA-limited chemostats. The working volume of each chemostat was 300 mL. After batch mode, pumps were turned on such that the dilution rate was approximately 0.018/hr. Mean and median cell volume and cell number were assayed using the Coulter counter. Each chemostat was sampled 4x each for metabolites after Coulter counter readings and media pH stabilized.

### Metabolite sampling and normalization

#### Metabolite pool size sampling from chemostats

Metabolites were sampled according to the procedure described in Crutchfield et al. (81). 5 mL of chemostat culture was filtered and metabolism was immediately quenched using 1.5 mL of −20°C 40:40:20 ACN:MeOH:H_2_O. Samples were then concentrated by drying with nitrogen gas and subsequent resuspension in 100% HPLC water. These samples were then analyzed via reversed phase ion-pairing liquid chromatography coupled to a Thermo Fisher Scientific Exactive instrument with high mass accuracy, allowing untargeted analysis (82, 83). Samples were collected in quadruplicate. Media filtrate samples were analyzed using two triple-quadrupole mass spectrometers, one running in positive mode (Finnigan TWQ Quantum Ultra) that was coupled to HILIC chromatography (84) and one in negative mode (TSQ Quantum Discovery Max) that was coupled to reversed-phase ion-pairing liquid chromatography (85) as previously described (66).

Data were normalized by total cell volume as per Boer et al., 2010 (66) and then log-transformed. In Figures 4 and 5, each sample from a deletion strain was compared to the corresponding wild-type sample run immediately preceding. In Supplemental Figure S6, the data were similarly normalized for run order (which was not confounded with either strain background or nutrient limitation): briefly, a linear model ***Y*** *= a****X*** *+ b + ε* was fit to each metabolite vector **Y**, using run order as the regressor **X**. The residuals *ε* were then kept and visualized as a heat map, after subtracting out the average levels of metabolites in wild-type under glucose-limitation.

#### Metabolite labeling experiments

Metabolites were labeled by transferring cultures into media with either U-^13^C glycerol and U-^12^C ethanol, or 2- ^13^C EtOH and U-^12^C glycerol. Labeled substrates were provided by Cambridge Isotopes. After 8 hours of growth in the labeled medium, metabolism was quenched, and extracts were then concentrated 3x and analyzed with LC/MS as above. Three biological replicates were sampled per strain and condition. Isotope labeling patterns were corrected for natural abundance and impurity of the tracer (∼1% ^12^C) using least-squares.

## Data Availability

Source code used to perform the analyses is available from http://www.bitbucket.com/pbradz/isozymes. Steady-state metabolite ion counts are provided in Supplemental Datasets S1 and S2.

## Acknowledgements

The authors would like to thank Amy Caudy for providing strains and expertise, David Hess for providing strains, Jing Fan for code to correct isotopomer distributions, and Yifan Xu for helpful discussions and for providing the *cdc19-*E392A strain. This research was made possible by funding from the NIH (R01 GM-071966 to OGT), NSF (MCB-0643859 and CBET-0941143 to JDR), AFOSR (FA9550-09-1-0580 to JDR) and the DOE (DE-SC0002077).

## Author contributions

PHB, DB, OGT, and JDR designed the research plan; PHB performed the computational analysis, *aco2Δ* and *pyk2Δ* growth curves, and metabolite labeling and quantification experiments; PHB and PAG performed chemostat cultures; PAG constructed the *pyk2*Δ rescue strains and assayed their growth; and PHB and JDR wrote the manuscript. All authors read and commented on the manuscript.

## Note S1

### Key differences from previous analyses of expression data

Previously used statistics for assessing differential isozyme regulation include compendium-wide correlation (1) and PCoR (partial co-regulation, or the standard deviation of per-experiment correlation statistics) (2). One potential shortcoming of both compendium-wide correlation and PCoR is that they can be biased by the composition of the compendium itself, which, despite its diversity, is itself a highly biased sampling of experimental conditions and transcriptional states (3). If a given isozyme pair were strongly anticorrelated in only a small fraction of conditions assayed in the compendium, both PCoR and especially the overall correlation would tend to be dominated by the remaining conditions, lowering power.

Second, calculating a single overall correlation as in Ihmels et al. (1) also requires gene expression data to be concatenated into a single matrix. This requires normalization to minimize the contribution of technical variation between experiments; however, with hundreds of experiments spanning both single- and dual-channel microarrays, it is unclear how such normalization would be performed. Correlation across an entire compendium has also been shown to compare unfavorably to weighted per-dataset correlations in the context of function prediction (4).

Third, neither PCoR nor the overall correlation provide any condition-specific information, whereas the main purpose of the method we use is to associate the differential expression of isozymes with specific environmental perturbations. Finally, previous methods have not used thresholds derived from statistical hypothesis testing with false-discovery rate correction, which becomes particularly important in a compendium with hundreds of experiments (5), and all of these previous approaches used both positive and negative correlation, making them potentially sensitive to cross-hybridization (6, 7).

In our analysis, we took a different approach in which, within each experiment in our compendium, we identified statistically significant negative correlations between isozymes, and then looked for trends within and across clusters of related experiments.

## Note S2: Supplemental Methods

### Assembly of microarray compendium and calculation of normalized correlations

The datasets constituting the microarray compendium were drawn from two sources. First, we included 129 datasets previously collected by Hibbs et al. (4). Second, the Gene Expression Omnibus (8) was queried for series uploaded between January 1, 2005 and April 1, 2009 that included array data from *Saccharomyces cerevisiae* and contained at least 6 samples, returning 417 datasets. These 417 were then processed according to the following schema: (i) tabular files were extracted from the SOFT files; (ii) larger datasets were broken into logical datasets as per Hibbs et al. (4); (iii) redundant datasets (e.g., supersets of other logical datasets, or duplicates of the data collected by Hibbs et al.) were excluded; (iv) missing values were imputed via the KNNimpute tool in the Sleipnir library (9, 10) with default parameters; and (v) multiple probes for the same gene were collapsed into per-gene expression profiles using a maximum likelihood method (4). In total, 285 non-redundant expression datasets were added to the compendium from GEO.

Normalized correlations were calculated by taking pairwise Pearson correlations between all pairs of genes *x* and *y* in every dataset *d* as follows:

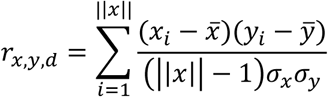

Here, *x* and *y* are the gene expression vectors in dataset *d*, *||x||* is the length of x, *x*_*i*_ and *y*_*i*_ are the values of each vector at index (i.e., array) *i*, and *σ*_*x*_ and *σ*_*y*_ are the standard deviations of *x* and *y*. Pearson correlations were then transformed according to the Fisher’s z-transform (i.e., hyperbolic arctangent).

-1

These scores were converted to standardized *z*-scores by dividing by the standard error 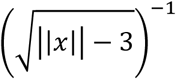. The *z-*scores were then used as the test statistic for a Wald test, yielding *p*-values. Finally, the *p-*values were corrected for multiple testing by conversion to *q-*values (5, 11).

### Logistic regression classification

We classified isozyme pairs into two groups based on whether they appeared, based on analysis of expression in our expression compendium, more like members of the same protein complex or more like random pairs. To do this, we took the binary differential expression vectors (see above) for pairs in the same protein complex and for random pairs, and fit a generalized linear model (using the “glm” function in R (12))) to classify pairs as one or the other:

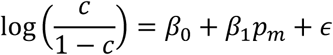

Here, *c* is the probability of a gene pair belonging to the same complex (vs. random pairs), *β*_0_ and *β*_1_ are parameters learned by the model, *p*_*m*_ is as above, and *ε* represents residual error. A value of *c* above 0.5 indicated that the pair was, at least weakly, more likely to be a “same complex” pair, and a value of *c* above 5/6 or below 1/6 was taken as a confident classification of a pair into the “same complex” or “random” categories. A graphical illustration is presented in Supplemental Figure 1.

### PAM clustering

Partitioning around medoids (PAM) was performed using the cluster package in R (13) with *k* = 3 based on a dissimilarity matrix constructed using the Jaccard distance.

## Supplemental Tables

**Supplemental Table S1.** Enrichments of metabolic pathways for reactions catalyzed by isozymes. Pathways were drawn from the Yeast Pathway database (14). *p-*values are from two-tailed Fisher exact tests, Bonferroni-Holm corrected for multiple testing. Pathway sets that were significantly enriched for isozymes at *p < 0.05* are shown in bold.

| Pathway | Isozymes | Other | *p*-value |
| --- | --- | --- | --- |
| Amino Acid Biosynthesis | 5 | 90 | 1 |
| **Glycolysis, Gluconeogenesis, Fermentation** | **9** | **12** | **5.44x10<sup>-8</sup>** |
| Lipid and Sterol Biosynthesis | 1 | 29 | 1 |
| NAD<sup>+</sup> Biosynthesis | 2 | 16 | 1 |
| Nucleotide Biosynthesis | 5 | 42 | 1 |
| **Pentose Phosphate Pathway** | **3** | **5** | **0.0245** |
| SAM Cycle | 1 | 2 | 0.86 |
| Storage Carbohydrates | 2 | 10 | 0.86 |
| **TCA and Glyoxylate Cycle** | **4** | **8** | **0.0122** |
| Other | 7 | 271 |  |
| Total | 39 | 485 |  |

**Supplemental Table S2.** Ten datasets with the greatest number of anticorrelated isozyme pairs. Source of the dataset (NP = not published), brief description, and a list of the anticorrelated isozymes are provided. The majority of these datasets have clear connections to glucose signaling (Ras, PKA) and availability. (Note that the *sua*5Δ deletion lacks cytochrome c and cannot grow on respiratory media.)

| Dataset | Brief description | Anticorrelated isozymes |
| --- | --- | --- |
| Meng (NP) | *sua5Δ* deletion vs. wildtype | MET12/MET13, HXK1/HXK2, HXK2/GLK1, SAM2/SAM1, PDC6/PDC5, PDC6/PDC1, DLD1/DLD2, TDH1/TDH3, URA5/URA10, UTR1/YEF1, ACS1/ACS2, CDC19/PYK2, DAK2/DAK1, ENO1/ERR3, ENO2/ERR3, GND2/GND1, GPM1/GPM2, GPM2/GPM3, GPP2/GPP1, IMD2/IMD4, PFK26/PFK27, PGM1/PGM2, SOL3/SOL4, TAL1/NQM1 |
| Smith 2008 | Growth of segregants on ethanol and glucose | HXK1/HXK2, HXK2/GLK1, PDC6/PDC5, PDC6/PDC1, PYC1/PYC2, NMA2/NMA1, ACO1/ACO2, YIA6/YEA6, DAL7/MLS1, ENO1/ERR3, ENO2/ERR3, GND2/GND1, GPD1/GPD2, GPM1/GPM2, GPM2/GPM3, |
|  |  | INM1/INM2, MHT1/SAM4, PFK26/PFK27, PGM1/PGM2, SOL3/SOL4, TAL1/NQM1, URA7/URA8 |
| Rossouw 2008 | Profiling of wine strains during fermentation | HXK1/HXK2, HXK2/GLK1, PDC6/PDC1, PDC5/PDC1, LYS21/LYS20, URA5/URA10, ACO1/ACO2, ACS1/ACS2, CDC19/PYK2, DAL7/MLS1, ENO1/ENO2, ENO2/ERR3, GDH3/GDH1, GND2/GND1, GPM1/GPM3, GPM2/GPM3, PGM1/PGM2, SOL3/SOL4, TAL1/NQM1 |
| Zhu 2009 | Heat shock | HXK1/HXK2, HXK2/GLK1, PDC6/PDC5, PDC6/PDC1, PYC1/PYC2, URA5/URA10, NMA2/NMA1, ACO1/ACO2, ACC1/HFA1, ACS1/ACS2, ASN1/ASN2, GDH3/GDH1, GPD1/GPD2, GPM1/GPM3, GPM2/GPM3, GPP2/GPP1, PFK26/PFK27, PGM1/PGM2, URA7/URA8 |
| Urban 2007 | Sch9/TORC1 signaling | NTH2/NTH1, PDC6/PDC5, PDC6/PDC1, PYC1/PYC2, ADE16/ADE17, UTR1/YEF1, ACC1/HFA1, ACS1/ACS2, DAK2/DAK1, DAL7/MLS1, ENO1/ERR3, ENO2/ERR3, GDH3/GDH1, GND2/GND1, HMG2/HMG1, PFK26/PFK27, SOL3/SOL4, TAL1/NQM1, URA7/URA8 |
| Guldal (NP) | Ras/PKA response (glucose regulation) | MET12/MET13, HXK1/HXK2, HXK2/GLK1, PYC1/PYC2, URA5/URA10, THI20/THI21, NMA2/NMA1, ACO1/ACO2, YIA6/YEA6, ACS1/ACS2, GDH3/GDH1, GPD1/GPD2, GPM1/GPM3, GPM2/GPM3, INM1/INM2, MHT1/SAM4, PFK26/PFK27, SOL3/SOL4, URA7/URA8 |
| Sill (NP) | Histone acetyltransferase study (ESA1) | PDC6/PDC5, URA5/URA10, NMA2/NMA1, CDC19/PYK2, DAK2/DAK1, ENO1/ERR3, ENO2/ERR3, GDH3/GDH1, GND2/GND1, GPM1/GPM2, GPM2/GPM3, IMD2/IMD3, IMD2/IMD4, INM1/INM2, MHT1/SAM4, PFK26/PFK27, PGM1/PGM2, SOL3/SOL4, TAL1/NQM1 |
| Singh 2005 | Dessication and rehydration on limiting glucose (lab strain, series 2) | MET12/MET13, SAM2/SAM1, PDC6/PDC5, PDC5/PDC1, PYC1/PYC2, URA5/URA10, NMA2/NMA1, ACS1/ACS2, CDC19/PYK2, GDH3/GDH1, GND2/GND1, GPM1/GPM2, HMG2/HMG1, INM1/INM2, PFK26/PFK27, TAL1/NQM1, THI21/THI22, URA7/URA8 |
| Sha 2013 | Oxidative stress | MET12/MET13, PDC6/PDC5, PDC6/PDC1, URA5/URA10, UTR1/YEF1, NMA2/NMA1, YIA6/YEA6, ACS1/ACS2, ASN1/ASN2, ENO1/ERR3, ENO2/ERR3, GND2/GND1, GPD1/GPD2, GPP2/GPP1, INM1/INM2, MHT1/SAM4, TAL1/NQM1, URA7/URA8 |
| Capaldi 2008 | *HOG1* mutants in either KCl or glucose upshift | HXK1/HXK2, HXK2/GLK1, PDC6/PDC5, PYC1/PYC2, URA5/URA10, ACO1/ACO2, ACS1/ACS2, GDH3/GDH1, GND2/GND1, GPD1/GPD2, GPM1/GPM3, HMG2/HMG1, INM1/INM2, PFK26/PFK27, PGM1/PGM2, SOL3/SOL4, URA7/URA8 |

**Supplemental Figure S1.**
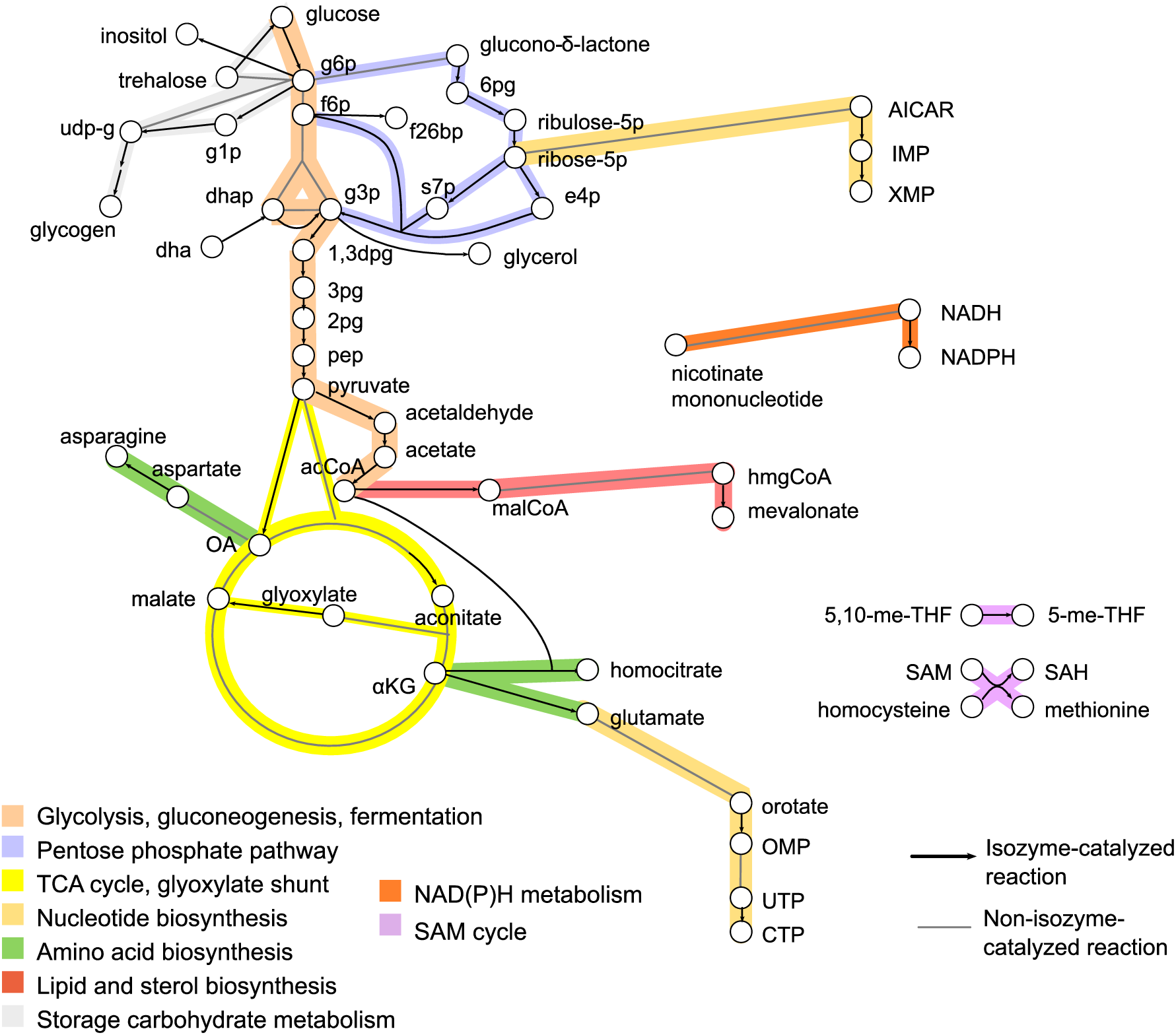
Co-localized metabolic isozymes in *Saccharomyces cerevisiae* are enriched in central carbon metabolism. Reactions catalyzed by metabolic isozymes of the same compartment are shown in black arrows; metabolites that are substrates or products of these isozyme-catalyzed reactions are shown as white circles. Other reactions are shown with a gray dotted line. Reactions are highlighted according to pathway (inset). While a large proportion of reactions catalyzed by these isozymes are in central carbon metabolism (glycolysis and gluconeogenesis, the TCA cycle, and the pentose phosphate pathway), comparatively few are in, for example, amino acid biosynthesis (green). Enrichment *p*-values are given in Supplemental Table S1.

**Supplemental Figure S2.**
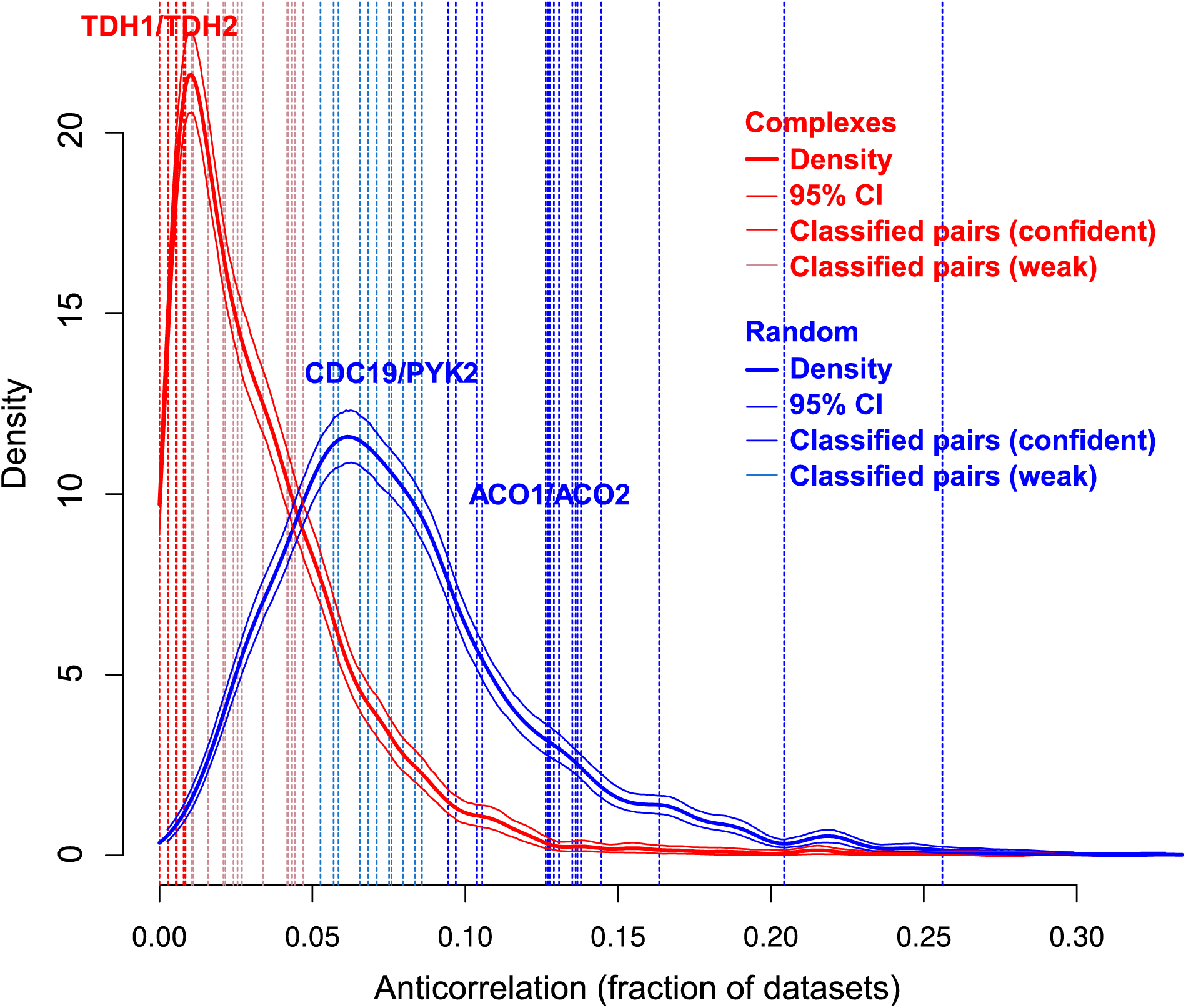
Illustration of logistic-regression-based classification of isozyme pairs. This classification relies on the average anticorrelation as defined in Methods under “Comparison of isozymes with other types of proteins.” The estimated distributions of average anticorrelation are plotted for pairs in the same protein complex (red) and random pairs (blue), with 95% confidence intervals plotted in light red and light blue. Blue and red dashed vertical lines represent individual isozyme pairs. Light red isozymes were classified as being more like “complex” pairs (P(*C*) > 0.5), with dark red isozymes classified particularly strongly as “complex” pairs (P(*C*) > 0.83). Light blue and dark blue isozymes were the same for “random” pairs (P(*C*) < 0.5 or P(*C*) < 0.17). Selected isozyme pairs are labeled in red or blue.

**Supplemental Figure S3.**
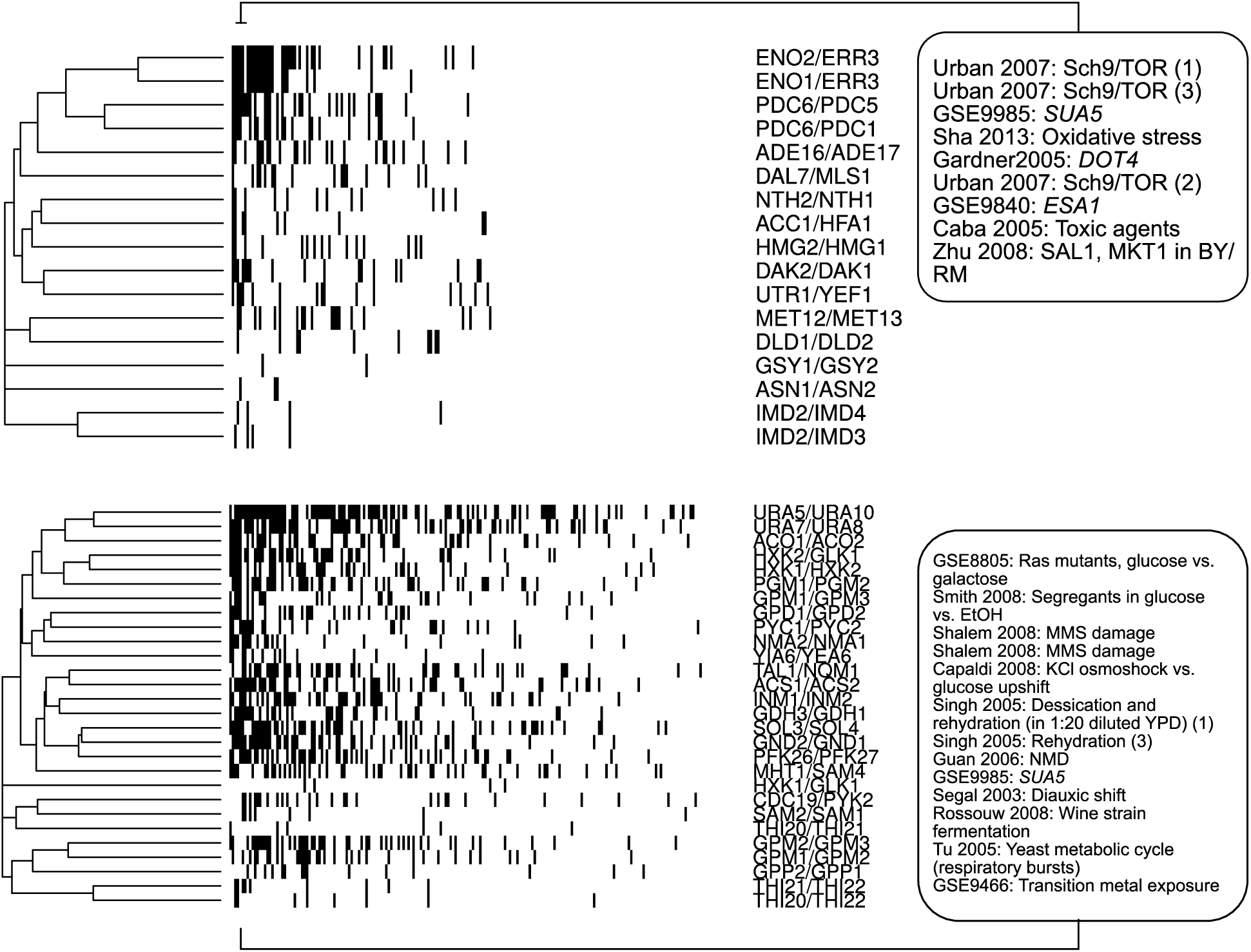
Partitioning around medoids (PAM) clustering of the binary differential expression matrix described in Methods (4.2) with three clusters revealed two coherent clusters, shown here. Black cells indicate that a given gene pair (row) was significantly anticorrelated (q < 0.1) in a particular dataset (column). Within each cluster, columns are sorted from most to least anticorrelation of isozyme pairs. The columns with the most anticorrelation of isozyme pairs for each cluster are highlighted on the right hand side of the figure. These datasets support a role for these isozyme pairs in the response to availability of glucose vs. other carbon sources.

**Supplemental Figure S4.**
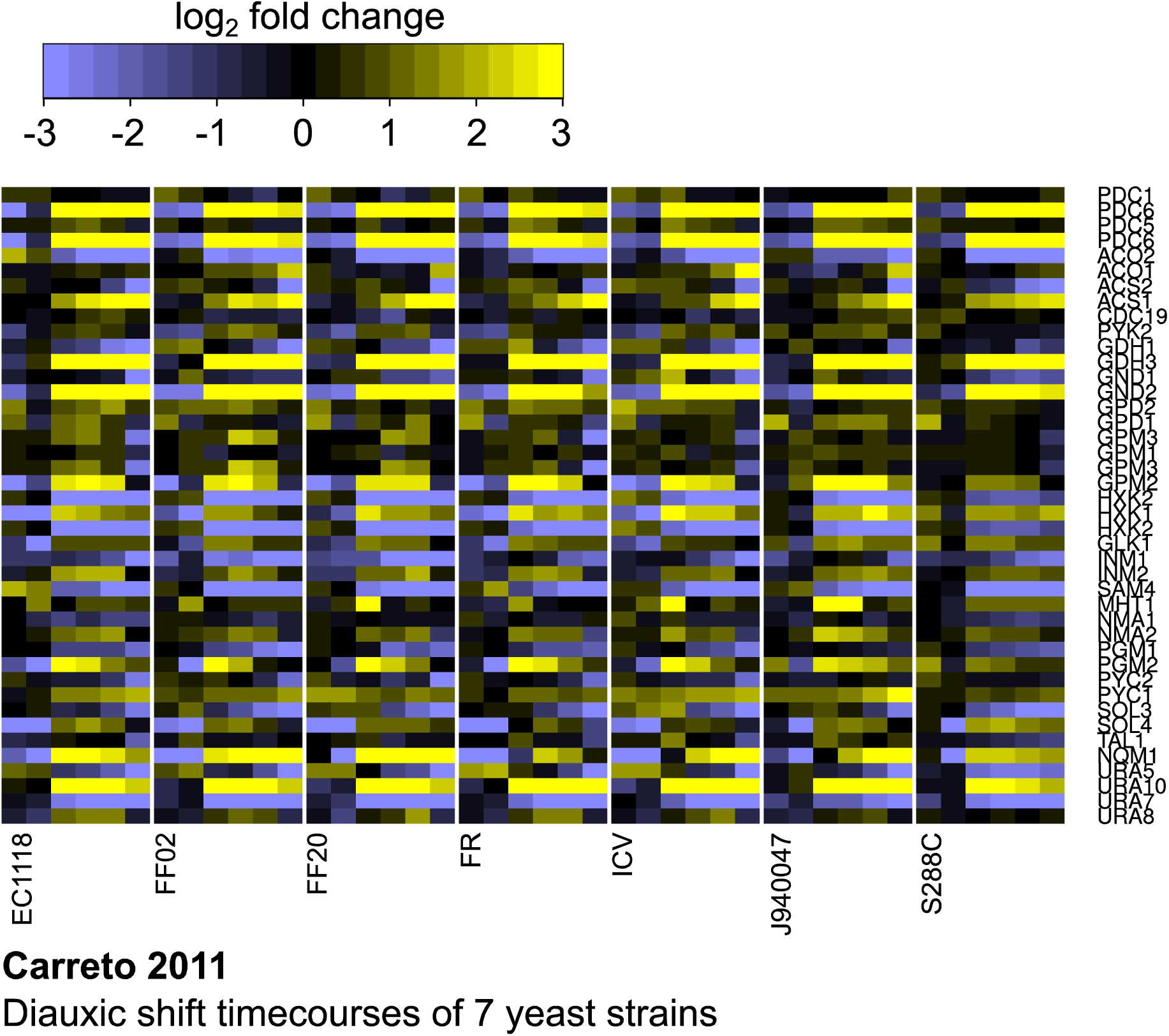
Anticorrelated expression of isozymes is conserved across diverged strains of *Saccharomyces cerevisiae*. Expression profiles are from a series of diauxic shift experiments conducted in wine, lab, and natural isolate strains of *S. cerevisiae* (15). As in Figure 4, these experiments show induction of one member of the pair (yellow) and repression of the other (blue) across the diauxic shift. Intensity corresponds to fold change; genes are grouped into isozyme pairs.

**Supplemental Figure S5.**
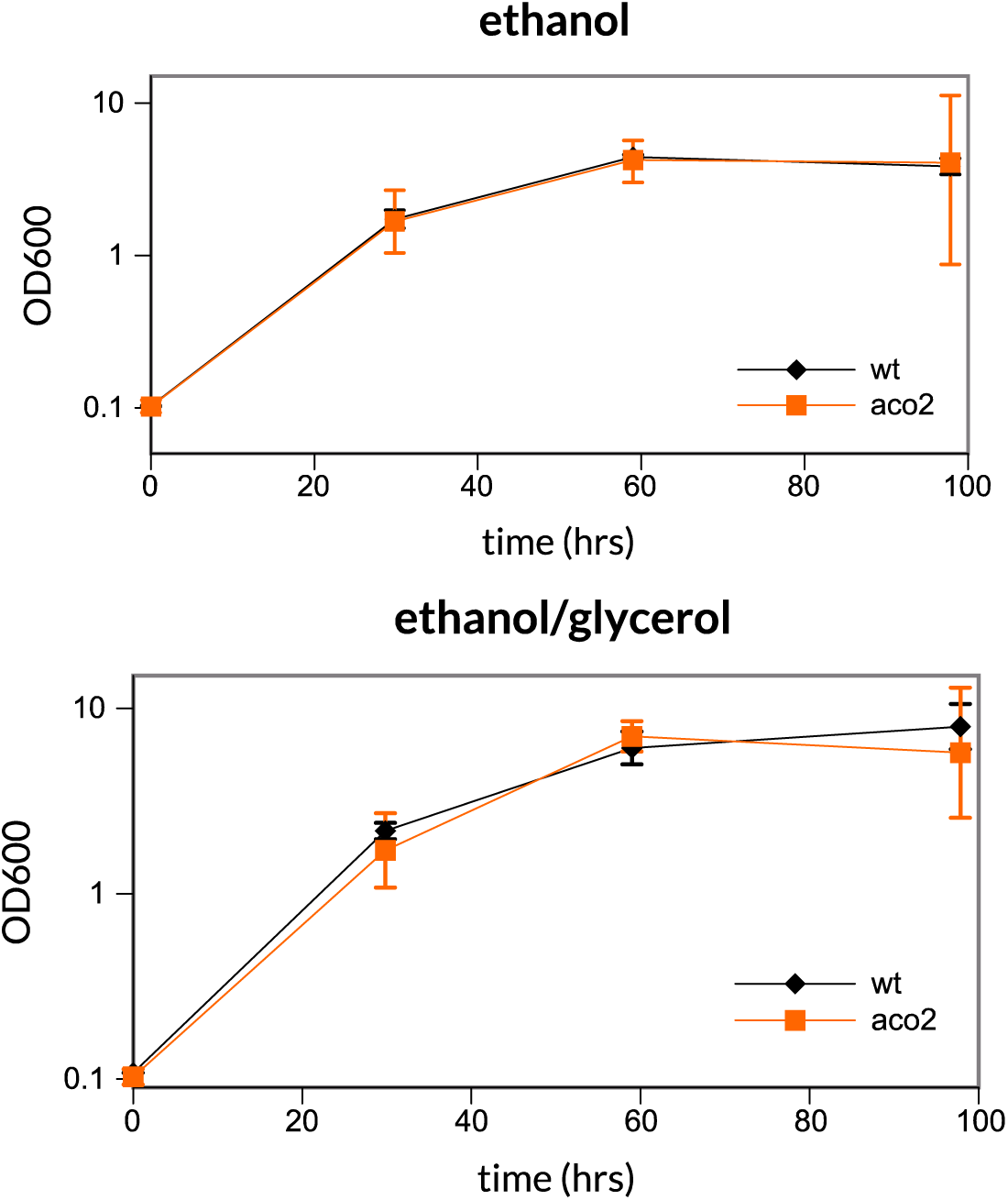
The aconitase 2 deletion *aco2*Δ has no growth defect on minimal media with a) ethanol or b) ethanol and glycerol as the carbon source. Compare growth on trehalose and glucose (Figure 4b). Error bars are 95% confidence intervals (n = 2 for wildtype; n = 4 for *aco2*Δ).

**Supplemental Figure S6.**
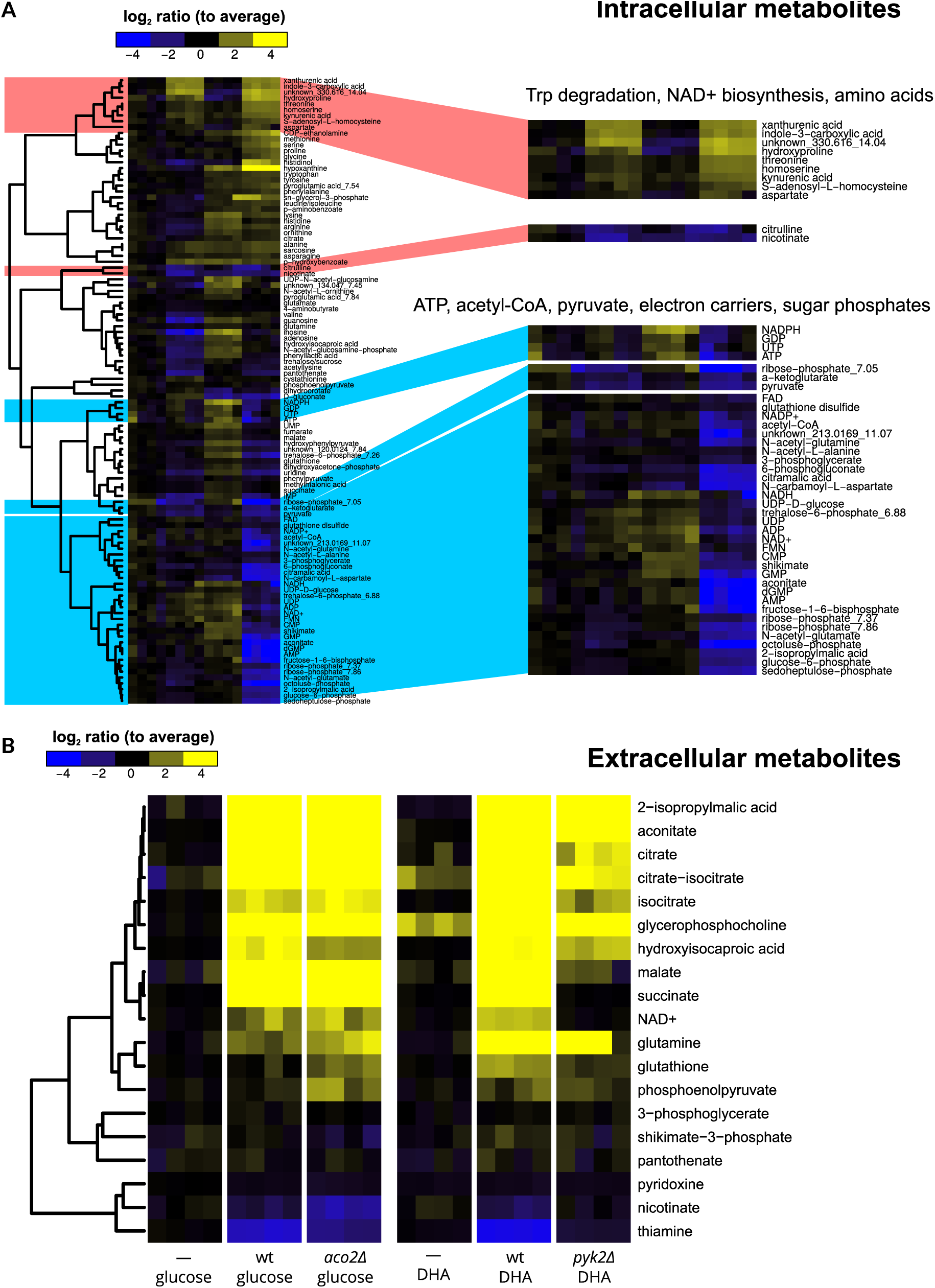
Intracellular and extracellular metabolic phenotypes of *aco2*Δ and *pyk2*Δ knockouts. Columns are repeated samples from single chemostats. Ion counts for intra- and extracellular metabolites are provided in Supplemental Datasets S1 and S2, respectively. a) Intracellular metabolomic profiles of glucose- or DHA-grown chemostat cultures of wildtype (wt), *aco2*Δ, and *pyk2*Δ strains. *pyk2*Δ strains show a decrease in glycolytic intermediates and phosphorylated compounds in general, but especially pyruvate and acetyl-CoA. A drop in ATP is also observed relative to wildtype (upper blue highlight). *aco2*Δ mutants show increases in tryptophan breakdown products (upper red highlight), and a drop in nicotinate (lower red highlight), indicating a shift from import of nicotinate to *de novo* biosynthesis of NAD^+^. b) Extracellular compounds from the same chemostat cultures. First and fourth groups of columns indicate pre-run chemostat media. Wild-type cultures show substantial excretion of TCA cycle intermediates, glutamine, and adenosine, and show strong uptake of the vitamins nicotinate and thiamine. Compared to wildtype, *aco2*Δ shows less uptake of extracellular nicotinate (a precursor to NAD^+^) and greater utilization of thiamine (used chiefly to make acetyl-CoA from pyruvate), while *pyk2*Δ shows very little uptake of either. *pyk2*Δ also shows sharply reduced excretion of TCA cycle intermediates, and excretes adenine in place of adenosine, possibly indicating limitation for five-carbon sugars.

**Supplemental Figure S7.**
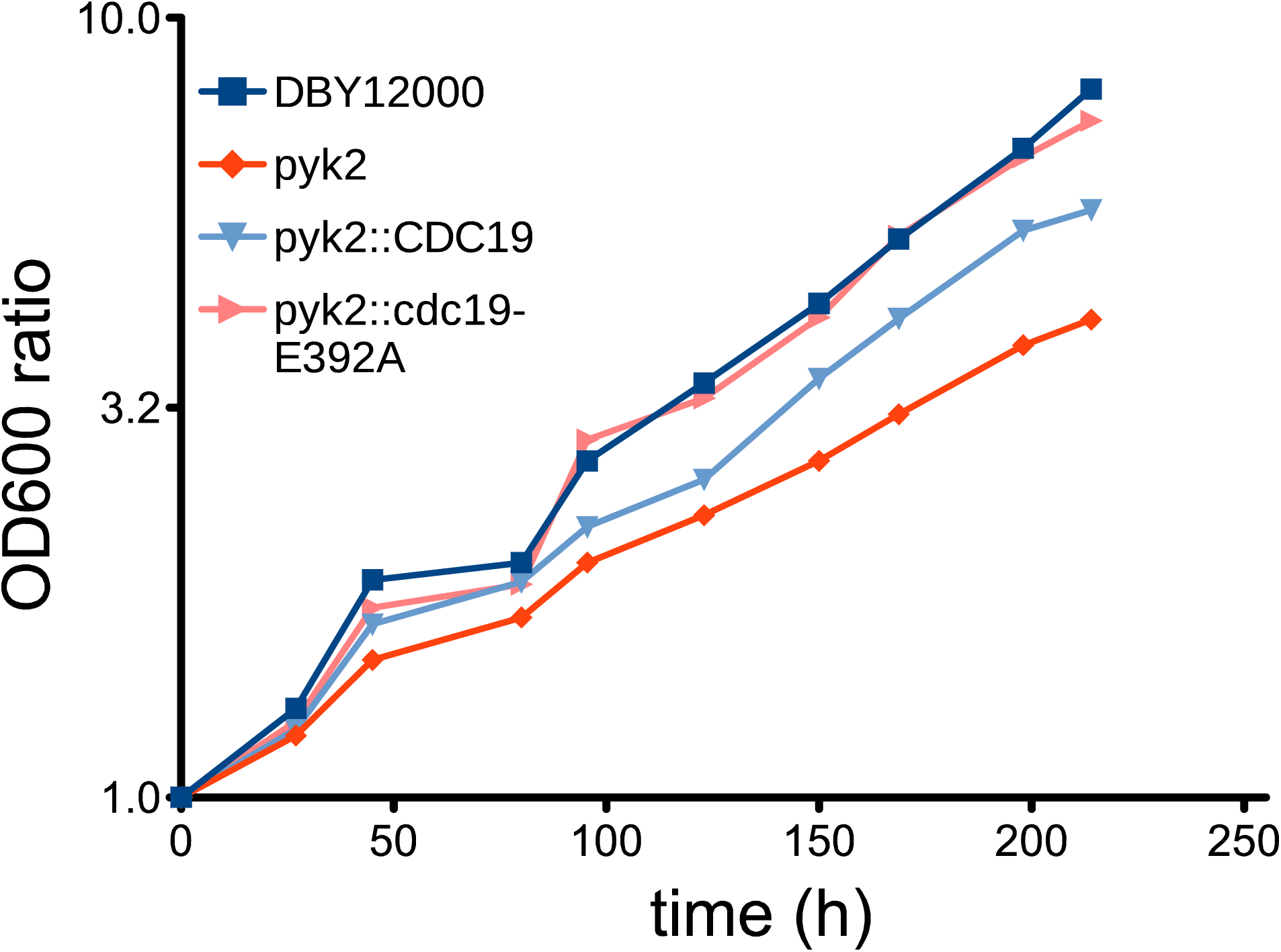
Growth of wt, *pyk2*Δ, *pyk2*Δ::*CDC19* and *pyk2*Δ::*CDC19E392A* rescue strains on dihydroxyacetone (n = 1) reveals rescue by *CDC19E392A*, a mutant of *CDC19* that is FBP-insensitive, but only incompletely by wild-type *CDC19*. Growth (y-axis) is expressed as a ratio of the OD600 at a given timepoint to the OD600 at time 0.

